# Adaptation of RiPCA for the Live-Cell Detection of mRNA-Protein Interactions

**DOI:** 10.1101/2023.09.07.556740

**Authors:** Dalia M. Soueid, Amanda L. Garner

## Abstract

RNA-binding proteins (RBPs) act as essential regulators of cell fate decisions through their ability to bind and regulate the activity of cellular RNAs. For protein-coding messenger RNAs (mRNAs), RBPs control the localization, stability, degradation, and ultimately translation of mRNAs to impact gene expression. Disruption of the vast network of mRNA-protein interactions has been implicated in many human diseases, and accordingly, targeting these interactions has surfaced as a new frontier in RNA-targeted drug discovery. To catalyze this new field, methods are needed to enable the detection and subsequent screening of mRNA-RBP interactions, particularly in live cells. Using our laboratory’s RNA-interaction with Protein-mediated Complementation Assay (RiPCA) technology, herein we describe its application to mRNA-protein interactions and present a guide for the development of future RiPCA assays for structurally diverse classes of mRNA-protein interactions.

## INTRODUCTION

It is well-established that protein synthesis is regulated by the dynamic networking of messenger RNAs (mRNAs) and proteins. Indeed, nearly all facets of an mRNA’s metabolism, its splicing, nucleocytoplasmic export, subcellular localization, functional engagement with the translation machinery, and mode and rate of degradation, are influenced by its cadre of bound RNA-binding proteins (RBPs) (Figure 1).^1^ Initial parameters used to define an RBP were based on the presence of one or more RNA-Binding Domain(s) (RBD), which recognize specific sequences or structural motifs within an RNA substrate.^2^ The most common RBDs include RNA-recognition motifs (RRMs),^3^ zinc-finger domains (ZnF),^4^ double-stranded RNA-binding domains (dsRBDs),^5^ and K-homology (KH) domains;^6^ however, others exist as well.^7^ Using this categorization, over 1,500 canonical RBPs have been identified, with ∼45% of those acting on mRNA targets;^7^ although non-canonical RBPs have also be reported and these numbers are likely to be much larger.^8^ Considering the significance of mRNA-protein interactions in ensuring maintenance of cellular homeostasis, dysregulation of this networking contributes to the development of several human diseases including cancers, neurodegenerative, cardiovascular, and autoimmune diseases.^9–14^ Accordingly, the targeting of RNA-protein interactions (RPIs) with small molecules has emerged as a new area of RNA-targeted drug discovery.^15, 16^

**Figure 1.**
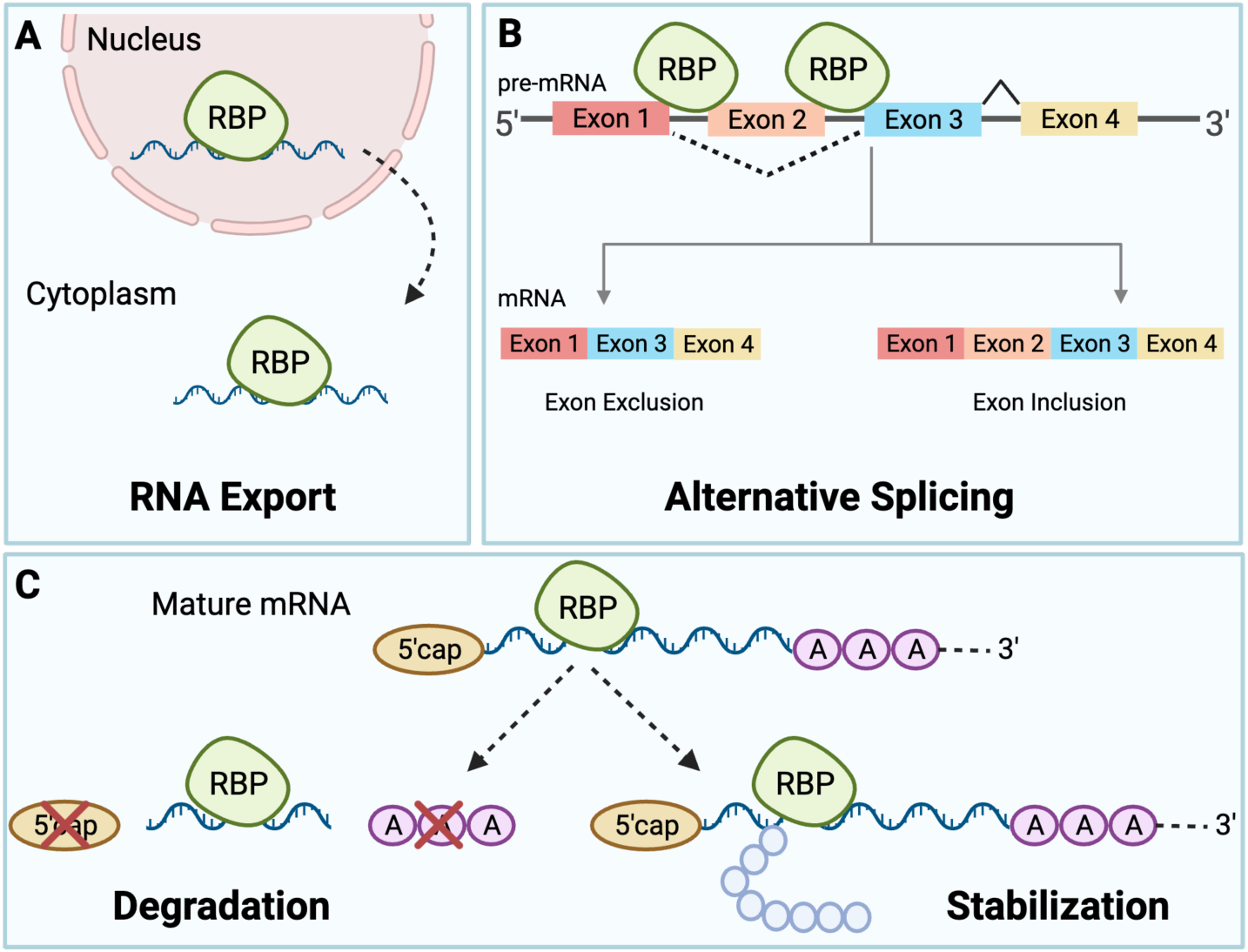
Cellular functions of mRNA-protein interactions. (A) RBPs regulate RNA localization, namely from their site of transcription in the nucleus to their site of translation in the cytoplasm. (B) RBPs serve as essential regulators of alternative splicing of pre-mRNA targets by binding to transcripts and facilitating exon exclusion or inclusion. (C) RBPs stabilize mRNAs to enhance translational efficiency or destabilize an mRNA transcript resulting in translation repression and/or degradation.

To enable these efforts, our laboratory has worked to develop cell-based assay technology for detecting RPIs, RNA-Interaction with Protein-mediated Complementation Assay (RiPCA) (Figure 2).^17–19^ RiPCA is an RPI complementation assay that utilizes Promega’s split nanoluciferase (NanoLuc^®^) technology where the nanoluciferase enzyme has been cut into small and large subunits, SmBiT and LgBiT, respectively.^20^ As these subunits have poor affinity for one another,^20^ this ensures that reconstitution of the enzyme, and subsequent signal generation, are driven by the affinity of the RPI. In brief, Flp-In^™^ HEK293 cells that stably express a SmBiT-HaloTag^21^ fusion protein are transiently co-transfected with an RBP-LgBiT plasmid and RNA substrate modified to bear a chloroalkane HaloTag^®^ ligand. HaloTag^®^ is a protein tag developed by Promega that can react with chloroalkane ligands;^21^ thus, by engineering SmBiT to bear HaloTag,^®^ the SmBiT-HaloTag fusion protein can be labeled by the RNA substrate in a biorthogonal manner in cells. Binding of the RBP to the RNA-SmBiT-HaloTag conjugate brings LgBiT in proximity to SmBiT, promoting reconstitution to form the full-length, catalytically active NanoLuc^®^ enzyme. Following treatment with NanoLuc^®^ substrate, chemiluminescence signal is produced to enable RPI detection. Using this technology, we have successfully detected RPIs between hairpin loop pre-microRNAs and their protein binding partners, including pre-let-7s with Lin28 and the interactions of Musashi1/2 with pre-let-7d and pre-miR-18a.^17, 19^ Yet, it was unclear if RiPCA, which is dependent upon proximity-mediated detection, would be applicable to more structurally complex RPIs. Demonstration of such applicability is critical as the development of a generalized method for RPI detection and screening would be highly advantageous for the field.

**Figure 2.**
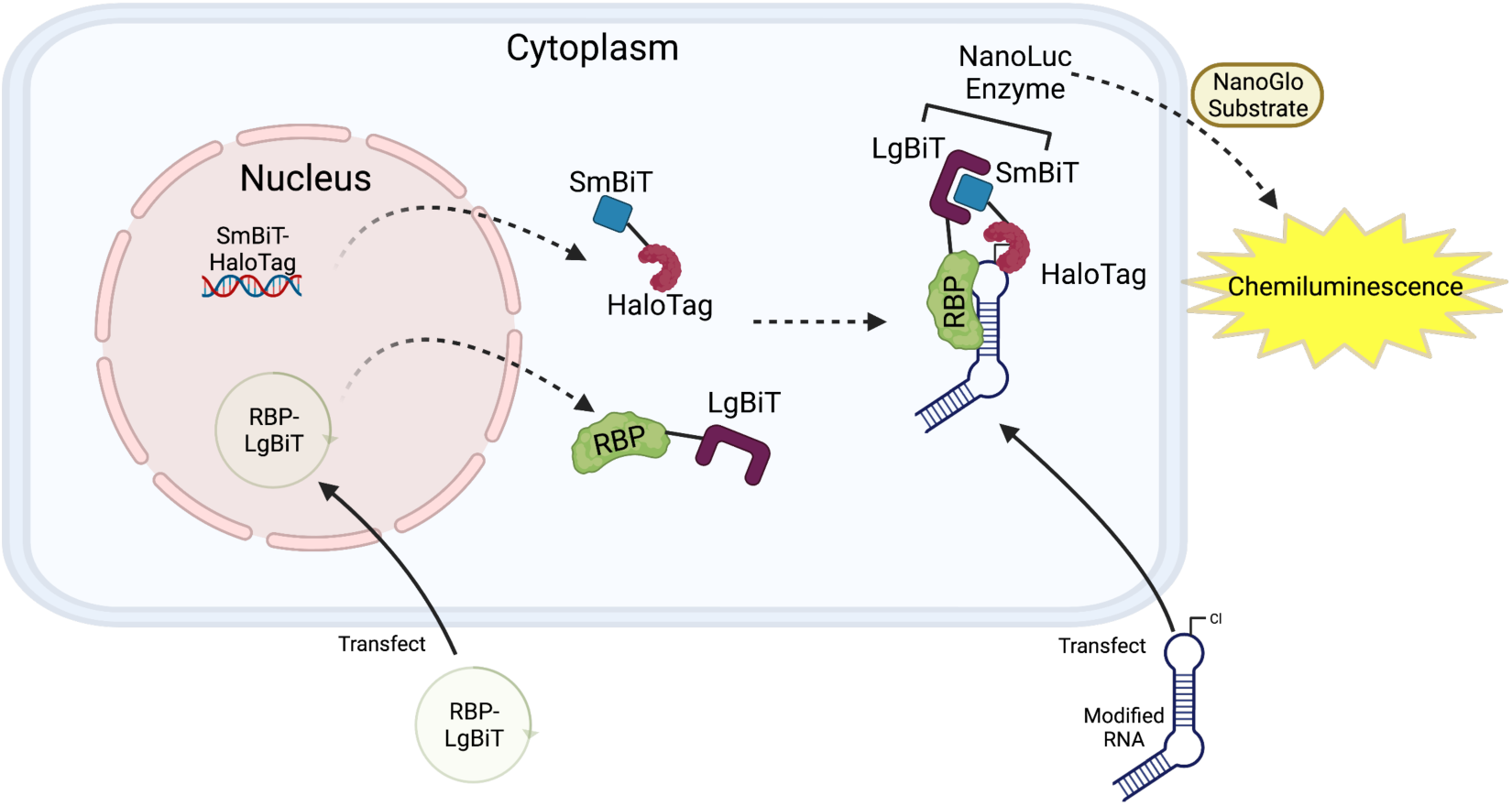
Schematic representation of RNA-interaction with Protein-mediated Complementation Assay (RiPCA). SmBiT = small subunit of nanoluciferase. LgBiT = large subunit of nanoluciferase.The chloro (Cl) modification depicted on the modified RNA implies a chloroalkane-tagged RNA substrate.

In addition to containing the code to make proteins, mRNAs contain regulatory regions upstream and downstream in the 5′ and 3′ untranslated regions (UTRs), respectively.^22^ While the 5′ UTR is predominately responsible for the regulation of translation initiation,^23^ the 3′ UTR is responsible for regulation of mRNA translation, as well as regulation of stability, localization, and polyadenylation.^24^ Translation initiation is driven in either a 5′ m^7^GpppX cap-dependent manner where the ribosome is recruited to the 5′ UTR by binding of RBPs to the cap structure to promote scanning towards the start codon,^25^ or a cap-independent mechanism where the ribosome is recruited to an internal ribosome entry site (IRES), which is a structural element within the 5′ UTR that can promote assembly of the translation machinery on the mRNA.^25^ The 3′ UTR, on the other hand, contains many structural elements which dictate mRNA fate and serve as hot spots for RBP binding.^26^ These *cis-*regulatory elements contain various sequence motifs which recruit *trans*-acting factors such as RBPs to carry out specific cellular functions. As more than half of human genes generate alternative 3′ UTRs, this serves an essential element governing gene expression, particularly at cell type- and tissue-specific levels.^26^

Current methods for detecting mRNA-protein interactions in live cells include fluorescence resonance transfer (FRET)- and trimolecular fluorescence complementation (TriFC)-based approaches.^27–30^ While these methods have been successful, particularly in the imaging of RPIs, there are several disadvantages of these technologies that we hypothesized that RiPCA could overcome. For example, FRET is highly dependent upon the stoichiometry of the donor- and acceptor-labeled biomolecules and typically requires high levels of expression,^31^ which may be toxic to cells depending on the RNA and RBP or present issues of non-specificity.^32^ For TriFC, while this approach offers advantages over FRET, such as enhanced sensitivity due to decreased background and less dependence upon the concentration of the interacting partners,^31^ a significant drawback is the pseudo-irreversibility of the reassembly of the split fluorophore, which is an advantage for detection of weak or transient interactions, but limits its use in measuring dynamics, such as inhibition by a small molecule.^30^

Given the prevalence of dysregulated mRNA-protein interactions in many human diseases, we sought to evaluate RiPCA’s ability to be adapted to studying these larger and more complex interactions. Focusing on three classes of mRNA 3′ UTR motifs, herein we report the first use of RiPCA for interactions beyond pre-miRNA-protein interactions. Moreover, we highlight challenges and lessons learned through our efforts in developing RiPCAs for several RPIs to guide others in the field who are interested in applying this technology to their RPI of interest.

## MATERIALS & METHODS

### Materials

Chemically synthesized RNAs (deprotected, desalted, and HPLC purified) containing a biotin at the 5′ end of the sequence and an internal 5-aminohexylacrylamino uridine (5-LC-N-U) modification were purchased from Horizon Discovery Biosciences and used as received for the labeling reaction. HaloTag^®^ Succinimidyl Ester (O2) Ligand was purchased from Promega (Cat #1691), dissolved in DMSO, and stored in single-use aliquots in amber tubes at -80 °C. The Nano-Glo^®^ Live Cell Assay System was purchased from Promega and used as received (Cat #N2012). TransIT-X2^®^ transfection reagent was purchased from Mirus (Cat #6000) and used as received.

### Cell culture

Flp-In^TM^-293 SmBiT-HaloTag or SmBiT-HaloTag-NLS cells, which express SmBiT-HT in the cytoplasm and nucleus, respectively,^17^ were cultured and maintained in DMEM (Corning Cat #10-017-CV) supplemented with 10% FBS (Atlanta Biologics S11550), L-glutamine (Gibco Cat #25030081), and 100 μg/mL hygromycin B (Gibco Cat #10687010) at 37 °C with 5% CO_2_ in a humidified incubator. Upon thawing, cells were passaged at least once before use in an experiment. Cells were passaged using trypsin-EDTA (0.25%) (Gibco Cat #25300054) ∼15−20× after thawing before returning to a new stock of cells.

### Cloning

Plasmids with RBP-LgBiT sequences were cloned using standard cloning techniques into a mammalian CMV-promoter containing pcDNA3 vector. DH5αF’IQ competent cells (Thermo Fisher cat. 18288019) were treated with Mix & Go! *E. coli* Transformation Kit (Zymo Research cat. #T3001) and used for cloning. Plasmid sequences can be found in the Supporting Information. Plasmids were prepared using a QIAprep Spin Miniprep Kit (QIAGEN cat. #27106) followed by ethanol precipitation for further purification. Purified plasmids were diluted to 3.9 ng/μL working concentrations and stored in 1.5-mL LoBind^®^ microcentrifuge tubes (Eppendorf cat. #022431021).

### RNA bioconjugation

The sequences of all RNAs purchased (with their modifications) can be found in Table S1. RNA substrates were conjugated to HaloTag^®^ Succinimidyl Ester (O2). Equal volumes of 1 mM RNA (in 100 mM phosphate buffer pH 8) were mixed with 10 mM HaloTag^®^ Succinimidyl Ester (O2) (in DMSO). The reaction proceeded at room temperature for 2 h. RNA was precipitated by the addition of 0.11× volume of 3.0 M sodium acetate and 4 equivalents of cold 200-proof ethanol and pelleted at 15,000 rpm for 45 min at 4 °C. The pellet was air dried and resuspended in 100 mM phosphate buffer (pH 8.0) to a concentration of 1.0 mM and stored at -80°C. Purified RNA was diluted to 25 μM for the working concentration using phosphate buffer (pH 8.0) in 1.5-mL LoBind^®^ microcentrifuge tubes and the concentration was verified by nanodrop.

### RiPCA

The RiPCA 2.0 protocol using TransIT-X2^®^ transfection reagent was previously reported.^19^ Briefly, solutions containing room temperature OptiMEM, DNA, RNA, and TransIT-X2^®^ (added in that order, volumes in Table S4) were mixed by quickly vortexing and briefly centrifuging prior to a ∼20 min incubation at room temperature while cells were harvested. Cells were harvested, counted, and prepared as a solution of 200,000 cells/mL. 300 μL of this solution was added to each tube containing OptiMEM, DNA, RNA, and TransIT-X2^®^ using a multichannel pipette. Each tube was mixed by pipetting up and down before plating 100 μL per well from each condition, in triplicate, in a white-bottom, tissue culture-treated 96-well plate (Corning cat. #3917). The plate was incubated in a humidified incubator (30 °C and 5% CO_2_) for 24 h. After 24 h, the media was removed from each well and replaced with 100 μL of room temperature phenol red-free OptiMEM (Thermo Fisher cat. #11058021) and treated with 25 μL of NanoGlo^®^ Live Cell Reagent diluted to 1:20 according to the manufacturer’s recommendation. All chemiluminescence data was collected immediately on a BioTek Cytation3 plate reader.

### Data analysis

All data was analyzed using GraphPad Prism version 9.5.1 for Mac OS X. Normalized chemiluminescence values shown were calculated by dividing the signal obtained for cells transfected with both the LgBiT-tagged RBP (1.0 ng/well) and respective RNA substrate (0.45 μL; 33 nM) over the signal obtained for cells transfected with the LgBiT-tagged RBP alone (1.0 ng/well) (average signal from triplicate wells). Wells containing cells transfected with RBP only measures the non-specific interaction of SmBiT-LgBiT correlated with the expression level of the RBP. Error bars represent standard deviation.

### Western Blot

Flp-In^TM^-293 cells stably expressing SmBiT-HaloTag or SmBiT-HaloTag-NLS were plated at a density of 150,000 cells/well in a clear, tissue culture treated 6-well plate. After 24 h, cells were transfected with a mixture containing OptiMEM (200 μL), pcDNA3 plasmid containing the sequence of interest (1 μg), and PEI transfection reagent (3 μL from a 1 mg/mL stock). Transfection components were added in the order listed, vortexed, spun down, incubated for 15 min, and then added to cells after changing the media. 24 h after adding the transfection components, cells were harvested (rinsed with PBS first) using 500 μL RIPA buffer containing protease inhibitor cocktail. Lysates were sonicated at 50% Amp for 6 s and then spun down at 15,000 rpm for 5 min. Total protein was quantified using BCA, and 7 μg of total protein was loaded onto a 4−20% Tris Glycine gel and resolved using 135 V for 90 min. The gel was transferred to a PVDF membrane in Towbin’s Buffer (25 mM Tris pH 8.3, 192 mM glycine, 20% *v*/*v* methanol). The membrane was blocked in 5% non-fat milk for 1 h at 25 °C and then incubated with anti-LgBiT primary antibody overnight at 4 °C and secondary antibody for 1 h at 25 °C. Mouse Anti-LgBiT monoclonal antibody was purchased from Promega (cat. #N7100), HRP-linked anti-mouse IgG was used as a secondary antibody and purchased from Cell Signaling Technology (7076S), and actin-HRP was purchased from Santa Cruz Biotechnology (#sc-47778) and used as a standard. Clarity Western ECL Substrate (BIO-RAD cat. #17050661) was used for detection and proteins were visualized using a Bio-Rad ChemiDoc imaging system. Western blot images are shown in Figures S1−S3.

## RESULTS AND DISCUSSION

Unlike pre-miRNAs, which are based upon a common stem-loop structure of ∼60−80 nts, allowing usage of full-length RNA substrates in RiPCA,^33^ mRNAs are large (∼1,000 nts) and structurally complex.^34^ However, mRNA-protein interactions typically encompass only a limited number of nucleotides due to the small footprint of RBDs (3−8 nts).^35^ Thus, we hypothesized that we could utilize smaller mRNA fragments as substrates in RiPCA, and through these assay development efforts, we would be able to explore how RNA structure affects detection in RiPCA and how best to build assays for RNAs that exist in structures aside from hairpin loops.

As model mRNA-protein interactions, we focused our assay design efforts on three well-validated systems representing RBP binding to mRNA 3′ UTR motifs: (1) (CUG)_n_ expanded repeat RNA binding to MBNL1, Stau1, and CELF1;^36–38^ (2) *numb1* mRNA binding to Musashi proteins;^39^ and (3) *c-fos* AU-rich element (ARE) binding to AUF1 (affinities of each interaction, or similar interactions, can be found in Table S3).^40^ These models were chosen as they represent unique structured RNA elements with validated binding mechanisms to their RBP binding partners. For each RNA substrate, secondary structures were predicted using the MC-Fold and MC-Sym pipeline (Figures 3A, 3D, and 3F)^41^ and 3D structures were predicted using the RNAComposer 3D modeling server^42, 43^ (Figures S5, S7, and S9) to identify uridine residues for modification to produce RNA substrates for RiPCA (Table S1).^17–19^ RNAs containing 5-aminohexylacrylamino-uridine (5-LC-N-U; structure in Table S2) modifications to replace select uridine residues and enable labeling with a chloroalkane HaloTag^®^ ligand (structures in Table S2) were designed to determine how the location of SmBiT-HaloTag labeling would affect detection in RiPCA. Based on the known RNA-binding sequences for each substrate, we also designed mutant RNA substrates to serve as negative controls for our assay development efforts (Figure 3 and Table S1).^39, 44–47^ For each RBP, both *N*- and *C*-terminal LgBiT tagging was explored (Figure 3).

**Figure 3.**
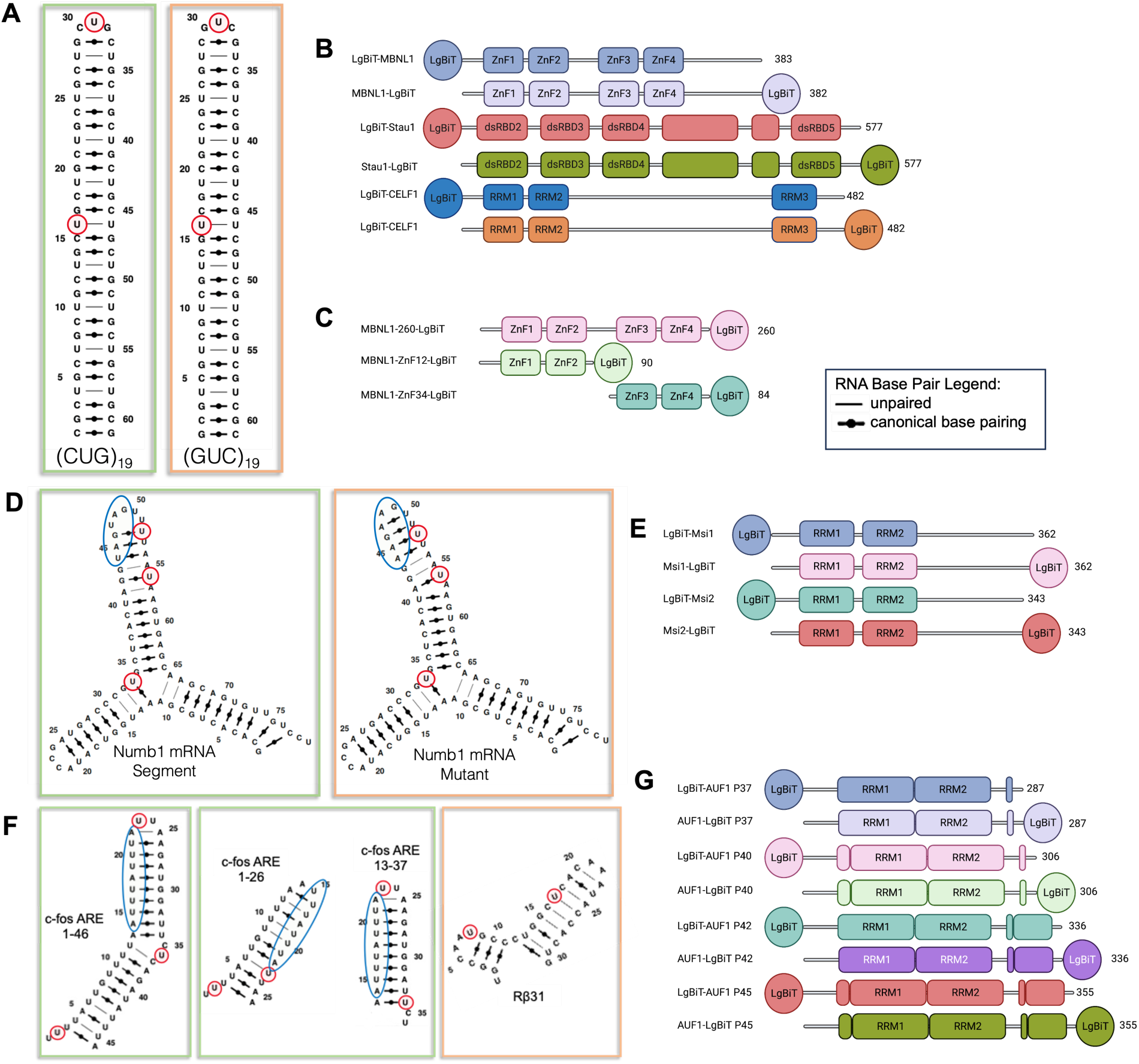
Model mRNA-protein interactions. Green boxes represent target RNA substrates. Orange boxes represent designed mutant RNA sequences. Blue circles indicate the RBP binding site on the RNA. Red circles indicate the site of 5-LC-N-U modification and HaloTag labeling. Legend for RNA base pairing is also depicted. (A) RNAs for the (CUG)_n_ expanded repeat systems. (B) LgBiT-RBP and RBP-LgBiT constructs tested with the (CUG)_n_ expanded repeat RNAs. (C) Truncated MBNL1-LgBiT constructs tested with the (CUG)_n_ expanded repeat RNAs. (D) RNAs comprising a motif from the *numb1* RNA and a corresponding mutant *numb1* motif. (E) LgBiT-Musashi and Musashi-LgBiT proteins tested with the *numb1* and *numb1* mutant RNAs. (F) RNAs comprising varying motifs from the *c-fos* ARE and a non-binding negative control, Rβ31. (G) LgBiT-AUF1 and AUF1-LgBiT proteins tested with *c-fos* ARE substrates depicted in (F).

### (CUG)_n_ Expanded Repeat RNA

Myotonic dystrophy type 1 (DM1) is a trinucleotide repeat disorder caused by a (CTG)_n_ expanded repeat in the *Dystrophia Myotonica Protein Kinase* (*DMPK*) gene, which is subsequently transcribed as an mRNA containing a (CUG)_n_ expanded repeat in the 3′ UTR.^48, 49^ DM1 Patients harbor repeats between 50 to 1,000 nts that form ultra-stable stem-loop structures^50^ which can bind and sequester the RBP Muscleblind-like 1 (MBNL1) in nuclear foci, disrupting its role as an alternative splicing regulator.^36, 44^ In addition to MBNL1, several other RBPs have also been demonstrated to bind to (CUG)_n_ expanded repeat RNA including Staufen1 (Stau1) and CUGBP ELAV like factor 1 (CELF1); yet, their role, if any, in DM1 is unclear.^37, 38, 51, 52^

As a model (CUG)_n_ RNA, we designed a (CUG)_19_ sequence^53^ containing an additional terminal GC base pair to stabilize the hairpin structure (Figures 3A and S5). 5-LC-N-U modifications were placed at U16 or U31 to probe SmBiT-HaloTag labeling of the stem and terminal loop of the RNA substrate, respectively (Figure 3A). As a non-binding negative control sequence, we similarly designed (GUC)_19_ RNAs,^44^ which maintain the same overall secondary structure, nucleobase composition, and modification sites as the binding sequence (Figures 3A and S5). Plasmids encoding *N*- and *C*-terminally tagged MBNL1, Stau1, and CELF1 (Figure 3B) were prepared, and protein expression was confirmed via Western blot (Figure S1). To determine the applicability of the (CUG)_n_ RPI in RiPCA, cells stably expressing SmBiT-HaloTag in the cytoplasm were transiently co-transfected with RBP-LgBiT plasmids (1 ng) and either U16 or U31 (CUG)_19_ RNA substrates (33 nM). Following 24 h incubation, cells were treated with NanoGlo^®^ Live Cell

Reagent and chemiluminescence signal was measured. For each mRNA-RBP pair, we calculated normalized chemiluminescence values by dividing the signal obtained for cells transfected with both an RBP plasmid and RNA substrate over the signal obtained by transfecting the RBP alone. Although LgBiT and SmBiT have poor affinity for one another (K_d_ of 190 μM),^20^ it is well established that expression level can influence background signal detection due to spontaneous formation of the LgBiT/SmBiT complex.^20^ By plotting calculated normalized chemiluminescence values for each assay, we are able to readily evaluate specificity of signal generated in RiPCA over background due to non-specific LgBiT-SmBiT binding.

As shown in Figure 4A, normalized chemiluminescence signal was calculated for all RBPs tested and negligible differences were observed between the U16 and U31 modified RNAs. While MBNL1 and Stau1 (both LgBiT orientations) exhibited strong chemiluminescence signal in RiPCA as expected, signal with CELF1 was much weaker. Upon closer inspection, the lower normalized chemiluminescence value calculated for LgBiT-CELF1 was not due to a lack of observed raw chemiluminescence signal, but rather resulted from higher RNA-independent background signal (Figure S4), reducing the assay window. We hypothesize that this could be due to the fact that CELF1 forms a homodimer in solution,^54^ allowing for a greater chance of LgBiT non-specifically interacting with SmBiT, a trend that we will revisit with another RBP that can multimerize in a similar manner. Alternatively, CELF1 binds only weakly to related (CUG)_n_ RNAs (Table S3). With respect to CELF1-LgBiT, this construct expressed poorly (Figure S1C), possibly due to the LgBiT tag interfering with the third RNA recognition motif (RRM) which facilitates dimerization of the RBP. Thus, in this case, the lack of signal observed was due to low raw signal obtained in RiPCA.

**Figure 4.**
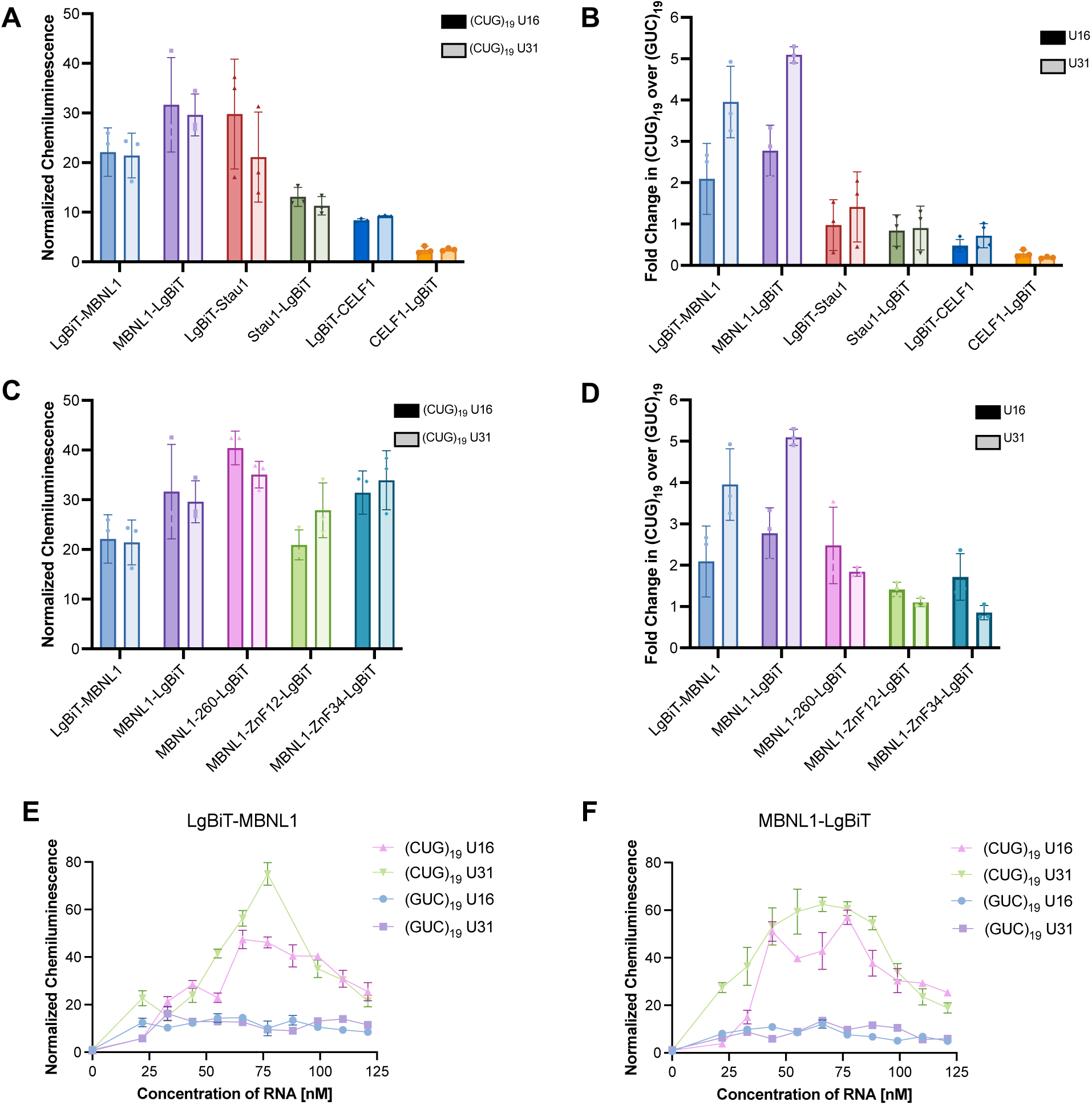
(CUG)_n_ expanded repeat RNA RiPCA. (A) Assay data with MBNL1, Stau1, and CELF1 in the cytoplasm. Normalized chemiluminescence values were calculated by dividing the signal obtained for cells transfected with both an RBP plasmid (1.0 ng/well) and RNA substrate (33 nM) over the signal obtained by transfecting the RBP alone. (B) Binding selectivity for (CUG)_19_ RNAs over non-binding (GUC)_19_ RNAs at each of the modification sites. (C) Normalized chemiluminescence values calculated in the cytoplasmic assay for various MBNL1 constructs. (D) Binding selectivity of truncated MBNL1 constructs at each of the modification sites. (E) and (F) RNA-dependence of (CUG)_n_ expanded repeat RNA RiPCA with MBNL1. Normalized chemiluminescence values calculated using (E) LgBiT-MBNL1 and (F) MBNL1-LgBiT (1.0 ng/well each). LgBiT-RBP = *N*-terminally tagged RBP. RBP-LgBiT = *C*-terminally tagged RBP.

We next investigated binding selectivity by performing RiPCA studies using the corresponding (GUC)_19_ RNA substrates. By comparing the fold-change in normalized chemiluminescence signals calculated for (CUG)_19_ RNAs over these non-binding sequences, we observed that only MBNL1 showed selectivity with a preference for the U31 modification site (Figure 4B). This selectivity is primarily driven by the lower signal obtained with the (GUC)_19_ U31 RNA in comparison to the (GUC)_19_ U16 RNA. As the interaction between MBNL1 and (CUG)_n_ repeat RNA is predominately driven by stacking of aromatic and arginine residues with the GC bases enabled via the bulges generated by the unpaired uridines in the stem of the RNA,^55^ and the U16 modification is located within the stem, this may result in a higher level of non-specific binding with the (GUC)_19_ U16 RNA.

To further probe the binding selectivity observed in RiPCA, we designed additional MBNL1 constructs. MBNL1 contains four ZnF domains; however, only ZnF2 and ZnF4 have been shown to directly interact with RNA.^56^ To probe the impact of ZnF1/ZnF2, ZnF3/ZnF4, as well as the *C*- terminal intrinsically disordered region (IDR), plasmids encoding the constructs shown in Figure 3C were prepared and tested in RiPCA with both (CUG)_19_ and (GUC)_19_ RNA substrates (Figure 4C). Removal of the IDR in MBNL1-260-LgBiT, and in turn placing LgBiT closer to the RNA-binding ZnF domain, resulted in higher signal than both full-length constructs (Figure 4C). The polypeptide linker between the tandem ZnF1/2 and ZnF3/4 has been shown to enable MBNL1 to adopt multiple conformations, and previous studies have demonstrated that MBNL1 containing all four ZnF domains can bind to two GC sequences separated by only one nt.^57^ We hypothesize that this explains the lower overall signal detected for the constructs containing only two ZnF domains due to reduction of binding of the protein and RNA to drive the interaction. As shown in Figure 4D, removal of the IDR or ZnF3/ZnF4 eliminated all preference for the (CUG)_19_ substrate. For the ZnF3/ZnF4 construct, while the highest level of selectivity was observed for the U16 modified (CUG)_19_ RNA, overall signal was low in comparison to the full-length MBNL1 constructs. Thus, using RiPCA, we conclude that while the full-length MBNL1 is not required to achieve high affinity, all domains are necessary to achieve specificity for (CUG)_n_ RNA.

As an additional characterization of binding selectivity, we performed dose response experiments by titrating in increasing amounts of the RNA substrates in RiPCA. Importantly, we observed a dose-dependent increase in signal for LgBiT-MBNL1 and MBNL1-LgBiT with the (CUG)_19_ substrates, while no change in chemiluminescence signal above background was detected with the non-binding (GUC)_19_ control RNAs (Figures 4E and 4F). Furthermore, a “hook effect,”^58^ was apparent at high RNA concentrations indicating saturation of the RNA and/or RBP without reconstitution of the split nanoluciferase enzyme. Interestingly, a similar effect was not observed with LgBiT-Stau1 which showed non-selective binding to both (CUG)_19_ and (GUC)_19_ RNAs (Figure S6). Combined, these results provide further confidence that the signal we observed in RiPCA was driven by the RBP binding to the RNA and further demonstrates the high preference of MBNL1 for (CUG)_n_ repeats in line with DM1 disease pathology.

### *Numb1* mRNA: 5′ **UAGUAG 3′ Motif**

The *numb1* mRNA codes for a protein which is important for controlling cell fate decisions in the peripheral and central nervous systems and acts as an inhibitor of Notch signaling.^59^ Accordingly, *numb1* is tightly regulated at the post-transcriptional, translational, and post-translational levels.^60,61^ The Musashi (Msi) family of proteins, composed of Msi1 and Msi2, are highly conserved translational regulators containing tandem RNA-recognition motifs (RRMs) which bind the 3′ UTR of target transcripts containing a 5′ UAGUAG 3′ motif.^39^ Translational repression of *numb1* by Msi1/2 serves to increase Notch signaling in leukemia and several cancer types and has established Msi1/2 as a therapeutic target.^62^ We chose to evaluate the interaction between Msi1/2 and the *numb1* mRNA in RiPCA as our second model system.

For these experiments, a 79-nt sequence predicted to form a structured RNA displaying the single-stranded 5′ UAGUAG 3′ Msi1/2 recognition motif (Figures 3D and S7) was utilized.^45^ 5-LC-N-U modifications were placed at U33, U52, or U56 to probe SmBiT-HaloTag labeling at varying distances from the RBP binding site. As a non-binding negative control, we designed a *numb1* mutant which replaced U44 and U46 with adenines. This change of only two bases modified the binding site to 5′ AAGAAG 3′ to increase base pairing in the stem of the RNA, thereby removing the single-stranded nature of the binding site to diminish Msi1/2 binding by removing the key UAG binding motif.^63^ Other features of the mutant RNA such as nt length, overall secondary structure, and HaloTag^®^ modification sites were maintained to ensure comparable transfection efficiencies of the two RNAs. Plasmids encoding *N*- and *C*- terminally tagged Msi1 and Msi2 (Figure 3E) were prepared, and protein expression was confirmed via Western blot (Figure S2). The interaction between Msi1/2 and *numb1* was tested in a similar manner in RiPCA as described for the (CUG)_n_ expanded repeat RNA systems utilizing 1 ng of plasmid and 33 nM of RNA. Due to the presence of a nuclear localization signal in Msi1/2, we evaluated binding of this system in both the cytoplasm and nucleus by using either the SmBiT-HaloTag or SmBiT-HaloTag-NLS^17^ cell lines for cytoplasmic and nuclear RPI detection, respectively.

Normalized chemiluminescence values calculated for the Msi1/2 LgBiT constructs and *numb1* in the cytoplasm and nucleus are reported in Figures 5A and S8A, respectively. Cytoplasmic signals were higher for all constructs than those observed in the nuclear assays. A prominent trend observed across all assays was that the highest signals occurred with the U52 modification site on the *numb1* RNA substrate. Notably, this site is directly adjacent to the RBP binding site on the RNA leading us to hypothesize that proximity of the HaloTag^®^ ligand site and the RBP binding site positively influences detectable signal in RiPCA for this system. While a negligible difference between Msi1 and Msi2 binding to the *numb1* RNA was observed, higher signal was measured for the LgBiT-Msi constructs, where LgBiT is in closer proximity to the RBDs in comparison to Msi-LgBiTs.

**Figure 5.**
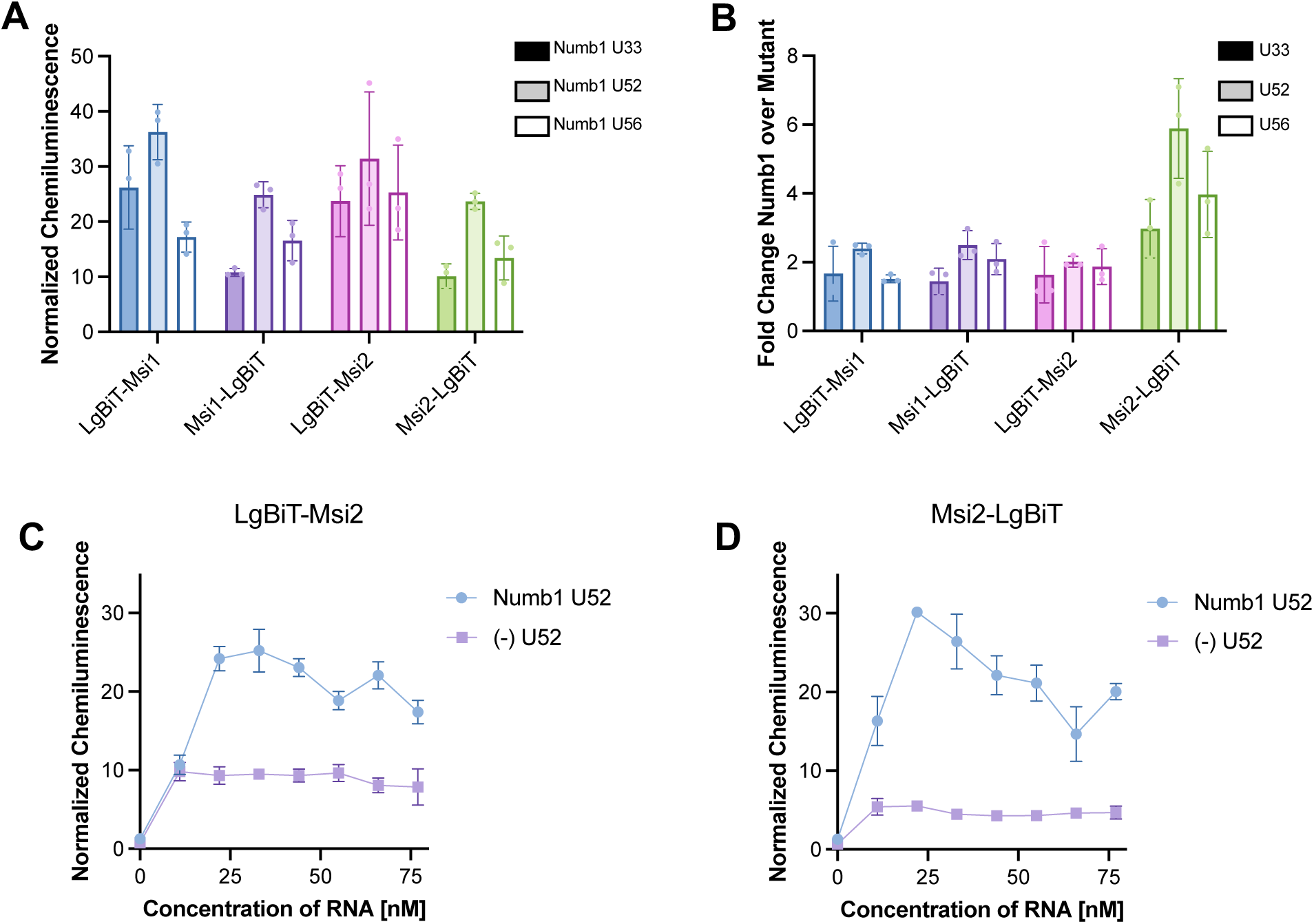
Numb1 RNA motif with Msi1 and Msi2 detected in the cytoplasm. (A) Normalized chemiluminescence values calculated in the cytoplasmic assay using 1.0 ng/well Msi1/2 constructs and 33 nM RNA. (B) Binding selectivity for *numb1* RNAs over non-binding mutant *numb1* RNAs at each of the modification sites. (C) and (D) RNA-dependence of Msi2 RiPCA. Normalized chemiluminescence values calculated using (C) LgBiT-Msi2 and (D) Msi2-LgBiT (1.0 ng/well each). LgBiT-RBP = *N*-terminally tagged RBP. RBP-LgBiT = *C*-terminally tagged RBP.

We next investigated binding selectivity of Msi1 and Msi2 by comparing the signals detected for the *numb1* and mutant *numb1* RNA substrates (Figure 3D). As shown in Figure 5B, the Msi2-LgBiT construct exhibited the largest preference for the target *numb1* RNA over the mutant RNA (cytoplasmic data shown; nuclear data in Figure S8B). The other Msi constructs displayed a specificity window of approximately 2-fold over the non-binding control. Interestingly, based on protein expression, Msi2-LgBiT levels were lower than that of LgBiT-Msi2 and the Msi1 constructs (Figure S2). Thus, we hypothesize that specificity of RiPCA signal is, in part, dependent upon expression of LgBiT-tagged RBPs. Further evidence supporting our hypothesis was obtained by performing dose-dependence assays. When titrating in increasing amounts of each RNA, we observed a dose-dependent increase in signal with the target *numb1* substrate with a consistent background for the mutant *numb1* RNA (Figures 5C and 5D), similar to our observations made with the (CUG)_n_ expanded repeat RNA and MBNL1. Because each RBP construct is normalized to its own background expression, the signals obtained for LgBiT-Msi2 and Msi2-LgBiT are comparable; however, a higher background measured with the mutant *numb1* substrate with LgBiT-Msi1 resulted in lower specificity of signal (Figures 5C and 5D). Thus, RBP expression must be controlled in RiPCA to ensure that strong signal window will be observed between binding and non-binding RNAs.

### *C-fos* mRNA: AU-Rich Element (ARE)

AU-rich elements (AREs) are one of the most common regulatory elements found within 3′ UTRs of mRNAs.^64^ AREs serve as a binding site for *trans*-acting factors such as RBPs, which influence the stability of the mRNA, its translation, and other mRNA fate decisions.^65^ Many of the transcripts which contain AREs encode tightly regulated factors such as inflammatory cytokines, growth factors, proto-oncogenes, and tumor suppressor proteins.^66, 67^ An RBP binding to an ARE can serve to stabilize or destabilize the RNA depending on the context of interaction.^66, 67^

To investigate this RPI system, we chose to use a 46-nt motif from the ARE found in the 3′ UTR of the *c-fos* mRNA as it is a well-established model for binding to the AU-rich element RNA-binding protein 1 (AUF1) also known as hnRNP D (Figures 3F and S9).^40, 67, 68^ AUF1 is expressed as four alternative splicing isoforms: the P40 and P45 isoforms contain the additional exon 2, the P42 and P45 isoforms contain the additional exon 7, and the P37 isoform is the smallest and does not contain exons 2 or 7 (Figure 3G).^69^ Absence or presence of these exons confers distinct biological properties and functions for each isoform. Inclusion of exon 7 affects nucleocytoplasmic transport,^70^ resulting in nuclear retention of the P42 and P45 isoforms.^71^ Exclusion of exon 2 is associated with a modest 3- to 5-fold higher affinity of the P37 and P42 isoforms for binding to ARE-containing RNAs;^72^ however, AUF1 oligomers, which are induced by RNA binding, are significantly more stable for the P42 and P45 isoforms.^72^

AUF1 binds to a tandem AUUUA pentamer within the *c-fos* ARE (Figure 3F), and similar to the *numb1* mRNA motif, we designed RNA substrates to examine internal modification sites at varying distances from the AUF1 binding motif.^67^ In addition to the 46-nt ARE RNA, we also designed two shorter substrates consisting of nts 1−26 and 13−37 (Figures 3F and S9), which differentially display the AUF1 binding motif. Each contained the same 5-LC-N-U modification sites as the full-length *c-fos* substrate for direct comparison. As a non-binding negative control, we employed the Rβ31 RNA previously shown to lack affinity for AUF1 P37 via an electrophoretic mobility shift assay (EMSA) (Figures 3F and S9).^46, 47^

Each alternative splicing isoform of AUF1 was cloned with an *N*-terminal and *C*-terminal LgBiT tag and tested at 1.0 ng/well of plasmid and a concentration of 33 nM for each RNA. While all AUF1 isoforms are able to shuttle between the nucleus and cytoplasm, P42 and P45 are predominately retained in the nucleus.^71, 73^ To enable direct comparison of the isoforms, all systems were tested in the SmBiT-HaloTag-NLS cell line for nuclear RPI detection. As shown in Figure 6A, normalized chemiluminescence values calculated for AUF1 with the *c-fos* ARE varied among each of the isoforms and between the *N*-terminal and *C*-terminal LgBiT constructs. As observed with Msi1/2, these differences are most likely due to the expression level of each of the fusion proteins (Figure S3). Also, in-line with data from previous RPI RiPCAs, we observed higher signal with the U2 and U23 modification sites, which lie on the same side of the hairpin as the AUF1 binding site (Figure 6A). We next sought to further characterize the *c-fos* ARE RiPCA using LgBiT-AUF1 P45 as a model as it had the highest normalized chemiluminescence values calculated and contains the additional alternative exons 2 and 7.

**Figure 6.**
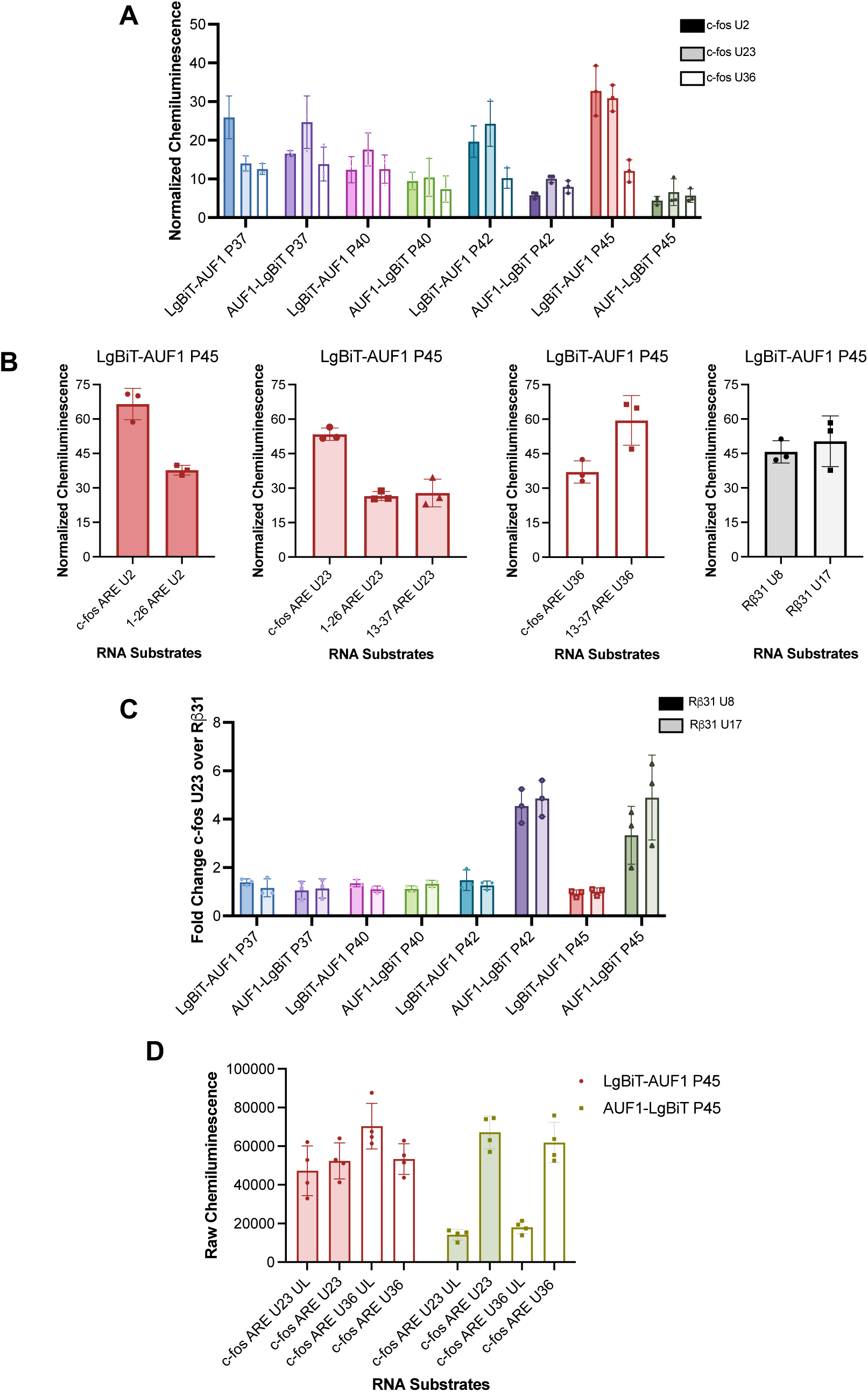
*c-fos* ARE RNA RiPCA. (A) Normalized chemiluminescence values calculated in the nuclear assay for all LgBiT-tagged AUF1 isoforms (1.0 ng/well) with 33 nM RNA. (B) Normalized chemiluminescence signals calculated with 0.5 ng/well of LgBiT-AUF1 P45 in the nucleus and all RNA substrates from Figure 3F (22 nM). Data is grouped by each of the RNA modification sites for direct comparison. (C) Binding selectivity for LgBiT-AUF1 and AUF1-LgBiT proteins for *c-fos* ARE U23 binding over Rβ31 U8 and U17. Values calculated using 1.0 ng/well of each plasmid with 33 nM RNA in the nuclear assay for the LgBiT-AUF1 proteins and the cytoplasmic assay for the AUF1-LgBiT proteins. (D) Raw chemiluminescence values of LgBiT-AUF1 P45 and AUF1-LgBiT P45 with *c-fos* ARE U23 and U36 with HaloTag^®^ ligand-modified and unlabeled (UL) RNAs. Values calculated using 1.0 ng/well of each plasmid with 33 nM RNA in the cytoplasmic assay.

AUF1 is a protein which exhibits a more complex binding mechanism to RNA than any of the previous RBPs tested using RiPCA. When binding to an RNA, AUF1 may bind as a dimer, trimer, or tetramer based on the length of the RNA substrate, and at least 34 nts are necessary for sequential dimer (tetramer) binding.^46, 47^ Based on previous reports, AUF1 can bind to RNA as a tetramer when an ARE is present in a sequence that is >30 nts. Thus, we anticipated that LgBiT-AUF1 P45 would bind to the 46-nt full-length *c-fos* substrate as a tetramer and the *c-fos* 1−26 and 13−37 truncated substrates as a dimer.

To investigate how multimerization affects detection in RiPCA, we tested LgBiT-AUF1 P45 across each of the RNA substrates shown in Figure 3F including the Rβ31 non-binding RNA control. As shown in Figure 6B, which displays normalized chemiluminescence values calculated for LgBiT-AUF1 P45 with all substrates, we observed various levels of signal generation for each of the full-length and truncated substrates. Unsurprisingly, the highest values calculated were associated with the *c-fos* ARE U2 and U23 substrates (Figure 6A). Interestingly, the truncated *c-fos* 13−37 produced equivalently high signal at the U36 site unlike the corresponding full-length U36 sequence (Figure 6B). We hypothesize that the U36 site in the *c-fos* 13-37 RNA may exhibit higher signal due to its structure. Upon examining predicted 3D RNA structures (Figure S9C), the helix in the *c-fos* 13-37 RNA is twisted, resulting in a conformation which brings the U36 site into an optimal position for LgBiT-AUF1 P45 to bind to the SmBiT-HaloTag. Moreover, we observed a 2- to 3-fold difference in normalized chemiluminescence between the full-length *c-fos* ARE RNAs at the U2 and U23 sites compared to the truncations with those same RNA modification positions (Figure 6B). This is in-line with the proposal that LgBiT-AUF1 P45 could be binding to the smaller substrates as a dimer, while it binds to the larger RNA as a tetramer^74^ or other higher-order oligomeric structure resulting in higher signal generation. The lower signal observed for the *c-fos* 1−26 substrate may indicate a preference for LgBiT-AUF1 P45 to bind to an RNA that is stabilized by formation of secondary structure which would further facilitate the binding of multiple subunits of the RBP.

Another notable feature observed from the LgBiT-AUF1 P45 assays with all nine of the RNA substrates was that there was only a small difference in signal generated with LgBiT-AUF1 P45 binding to the target *c-fos* ARE and Rβ31 negative control (Figure 6B). The Rβ31 RNA is also predicted to form a relatively stable hairpin structure (Figures 3F and S9D); thus, we hypothesize that this RNA structure could facilitate binding to LgBiT-AUF1 P45, resulting in signal detection. However, because previous *in vitro* studies utilized this RNA as a non-binding negative control with AUF1 P37,^46, 47^ there are likely other factors contributing to the signal detected in cells such as the expression of each of the AUF1 constructs and multimerization of the protein.

To investigate this further, we examined the signal obtained for each of the RBP-LgBiT plasmids transfected into SmBiT-HaloTag and SmBiT-HaloTag-NLS cells with no RNA present, i.e., the “background” signal obtained by the LgBiT and SmBiT subunits interacting non-specifically, driven by the expression level of the RBP (Figure S4). When comparing the signal generated by the AUF1 isoforms, notably LgBiT-AUF1 P45, we observed between 10−50× higher signal than any of the RBPs tested and ∼75× higher signal than a well which does not contain LgBiT-AUF1 P45 (Figure S4). Given the high expression of LgBiT-AUF1 P45, it is likely that we are detecting promiscuous binding events driven by overexpression. In-line with this, by analyzing the binding of LgBiT-AUF1 P45 to various RNA substrates beyond a single concentration, we observed comparable signal produced across all RNAs tested with no preference for *c-fos* ARE (Figure S10). Similar results were observed by altering the concentration of plasmid, and increasing the concentration of LgBiT-AUF1 P45 plasmid used yielded an increase in raw chemiluminescence signal measured and a corresponding decrease in normalized chemiluminescence calculated due to an amplification in background and reduction in the assay window (Figure S11).

From our analysis of background signal produced with the AUF1 constructs, we also found that the location of the LgBiT tag influenced the strength of the background signal produced with the AUF1 constructs in the nuclear vs. cytoplasmic cell lines. As shown in Figure S4, the *N*-terminally-tagged AUF1 proteins exhibited higher background signal in the nucleus, while the *C*-terminally-tagged AUF1 proteins exhibited higher background signal in the cytoplasm, indicative of high protein expression in each of these compartments as signal can only be generated by high local concentrations of both LgBiT and SmBiT components of the split nanoluciferase due to their poor inherent affinity for one another. As the nuclear import signal of AUF1 is located in a region near the *C*-terminus of the protein, we hypothesize that LgBiT at the *C*-terminus may hinder nuclear uptake. Indeed, there is no evidence in the literature of *C*-terminally tagged AUF1 proteins.

To determine how localization would affect specificity of detection for *c-fos* ARE, we evaluated the AUF1 constructs for binding to Rβ31 using the SmBiT-HaloTag cell line for cytoplasmic detection with the *C*-terminal LgBiT constructs and the SmBiT-HaloTag-NLS cell line for nuclear detection with the *N*-terminal LgBiT constructs (Figure 6C). Interestingly, only AUF1-LgBiT P42 and AUF1-LgBiT P45 displayed preference for the *c-fos* ARE over the Rβ31 RNA. Upon inspection of expression levels of these proteins, AUF1-LgBiT P42 and P45 are more poorly expressed than the other constructs and are nearer to endogenous protein levels, while the others are expressed to a higher extent (Figure S3). This data reiterates how expression level can play a role in the specificity of the signal generation.

Using LgBiT-AUF1 P45 and AUF1-LgBiT P45 as comparative models, we further investigated the specificity of signal generation. We performed experiments in the cytoplasmic assay using *c-fos* ARE U23 and U36 as RNA substrates. Additionally, we included *c-fos* ARE U23 and U36 which were not conjugated to the HaloTag^®^ ligand, and therefore, should not be able to be labeled by SmBiT-HaloTag in cells, referred to as “UL” (unlabeled) RNA. As previously reported^17^ and as demonstrated with the RBPs tested herein (Figure S12), signal detection in RiPCA is not achievable without the chloroalkane handle on the RNA. While the raw background signals for LgBiT-AUF1 P45 and AUF1-LgBiT P45 are similar in the cytoplasmic compartment, indicating similar expression levels of the protein, only LgBiT-AUF1 P45 generated signal in RiPCA with the UL RNA (Figure 6D). In addition to affecting the localization of the protein, we propose that the presence of the LgBiT tag at the *C*-terminus of the AUF1 protein may also affect its ability to multimerize and bind to RNA as a higher order structure. As mentioned previously, constructs containing the alternative exon 7 form more stable oligomers when bound to RNA. These results, notably, extend to non-substrate RNAs as well, and using the (CUG)_19_ RNAs from our MBNL1 RiPCA assays, we found that only AUF1-LgBiT P45 showed specificity across the different RNAs tested (Figure S13A). With LgBiT-AUF1 P45, some specificity could be detected, albeit only at low RNA concentrations (Figures S13B and S13C). Combined, these results demonstrate how overexpression of an RBP in RiPCA can result in lower signal detection and how each RPI system must be fine-tuned to prioritize a low background in combination with high raw signal.

### *Considerations* for Future RiPCA Systems

Given the data presented for the mRNA-protein interaction RiPCAs tested herein, as well as previous observations made with pre-microRNA-protein interactions RiPCAs,^17, 19^ we would like to conclude with lessons learned and provide considerations for setting up RiPCAs for additional RPIs.

#### RNA

For RNA substrate design, it is advisable to consider a length of RNA that may form a distinct secondary structure for enhanced stability. If the binding site of the RBP on the RNA is known, modification closer to the binding site should result in higher signal over background and over that generated using a non-binding control RNA.

#### RBP

For every plasmid that is cloned, is important to perform a test expression prior to testing in RiPCA to ensure that the construct expresses in the appropriate cell line. It is always useful to clone both an *N*-terminal and *C*-terminal LgBiT tag on the RBP-of-interest as it is difficult to predict which orientation will be optimal and, as demonstrated herein, signal in RiPCA is highly dependent on how well the protein construct expresses and how tolerable a tag is in a certain orientation. We have, however, noticed that a LgBiT tag that is in closer proximity to the RBD that binds to the RNA substrate is favorable for signal detection. Much larger proteins could possibly be truncated, as long as they are functional as individual domains, to increase the proximity of the LgBiT tag to the RBD; yet this could influence the specificity of the RBP as observed with MBNL1.

#### Assay considerations

Due to the low concentration of the plasmids and the small amount of reagent transfected per well in the assay, it is essential that each plasmid preparation is as clean as possible (added ethanol precipitation) to ensure that there is no variability between different plasmid preparations. It may be tempting to start with a higher concentration of the plasmid or RNA in the cells to ensure signal detection; however, normalized chemiluminescence values above background are only achieved when the background signal is low. We recommend starting with several low concentrations of plasmid (e.g., 0.5 ng, 1.0 ng, 1.5 ng) with a set concentration of RNA (25−75 nM) to determine an optimal assay window. From this point, a concentration of the plasmid can be set, and RNA can then be titrated to drive the signal and improve upon the assay window further, without contributing to the background. It is critical to balance the amount of the plasmid so that you have enough to detect signal, but not so much that your background is too high or that you begin to detect signal with off-target RNAs. Depending on the instrument used and the gain settings, the raw chemiluminescence values may fluctuate from day to day; however, the normalized values for an optimized system should remain consistent. Therefore, it is essential to include a “no RNA” negative control for each plasmid tested and for each concentration of the plasmid tested. Other aspects which may be altered to further optimize assay window include number of cells per well, amount of transfection reagent, and linker length of the HaloTag^®^ modification as previously described.^19^

## CONCLUSION

In conclusion, we have demonstrated the expanded use of RiPCA for RPIs occurring between RBPs and mRNA motifs. We have successfully been able to recapitulate the interactions occurring between various mRNA-protein pairs despite using smaller binding motifs of the target RNAs. Signal detection, in the cytoplasm or nucleus, was accomplished for each RBP which expresses in those cellular compartments. Further, we have been able to demonstrate selectivity in signal detection for target RNAs over similarly designed negative controls exemplifying our ability to distinguish between specific and non-specific signal detection. These experiments demonstrate promise in being able to use RiPCA more broadly to study a range of RPIs, including those occurring with various forms of non-coding RNAs and the RBPs that act on them. The largest hurdle to specific signal detection occurred when evaluating RBPs which bind as a dimer or tetramer to RNA. Coupled with overexpression of these protein, higher background values were observed, reducing the assay window. Further optimization of RiPCA to address reliance on RBP overexpression, as well as its adaptation for high-throughput screening, will be reported in due course.

## ASSOCIATED CONTENT

### Supporting Information

The Supporting Information is available free of charge at…

Cloning details, RNA probe sequences, supplemental figures and tables.

## ACKNOWLEDGEMENTS

This work was supported by the NIH (R01 GM135252 to A.L.G and T32 GM145304 to D.M.S). We would like to thank Dr. Sydney Rosenblum for cloning the Msi1/2 and CELF1 LgBiT plasmids. Figures 1, 2, 3B, 3C, 3E, and 3G and the Table of Contents were created with BioRender.com.

## ACCESSION IDs

MBNL1, Q9NR56; Stau1, O95793; CELF1, Q92879; Msi1, O43347; Msi2, Q96DH6; AUF1, Q14103

### ABBREVIATIONS

RBP: RNA-binding protein
RBD: RNA-binding domain
RPI: RNA-protein interaction
RiPCA: RNA interaction with Protein-mediated Complementation Assay
SmBiT: small subunit of nanoluciferase
LgBiT: large subunit of nanoluciferase
5-LC-N-U: 5-aminohexylacrylamino-uridine
UTR: untranslated region
MBNL1: Muscleblind-Like protein 1
Stau1: Stafen 1
CELF1: CUGBP ELAV Like Factor 1
Msi: Musashi
AUF1: AU-Rich binding Eactor 1
ARE: AU-Rich Element
IDR: Intrinsically Disordered Region
ZnF: zinc finger

## For Table of Contents use only

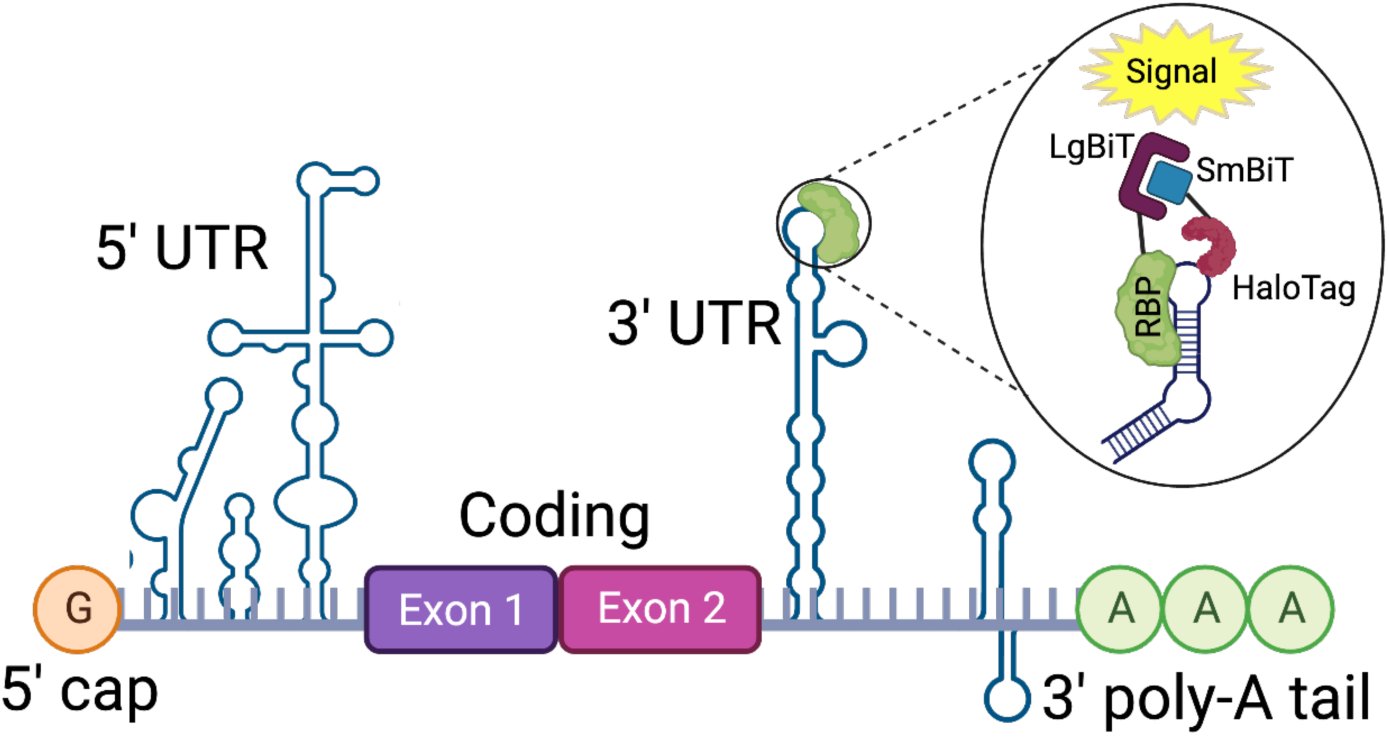

