## Supplemental Information for "Adaptation of RiPCA for the Live-Cell Detection of mRNA-Protein Interactions"

**A. Cloning** Pages S2−S15

**B. RNA Probes** Pages S16−S17

**C. Supplemental Figures** Pages S18−S24

**D. Supplemental Tables** Page S24

**E. References** Page S25

### A. Cloning

To clone all RBP constructs tagged with LgBiT, plasmids containing the LgBiT sequence were first cloned (referred to as *Plasmids A*, *B*, and *C*). Following this, RBP sequences were inserted to *Plasmids A*, *B*, and/or *C* to produce N- and C-terminally tagged RBPs.

*Plasmid A*: The LgBiT sequence was inserted into a pcDNA3 vector containing BamHI (fwd) and XhoI (rev) restriction enzyme sites. This plasmid was used to clone N-terminal LgBiT constructs (LgBiT-RBP). Cloning of this plasmid was previously reported.[1]

*Plasmid B*: The LgBiT sequence was inserted into a pcDNA3 vector containing HindIII and EcoRI restriction enzyme sites. This plasmid was used to clone N-terminal LgBiT constructs (LgBiT-RBP). Forward primer inserts Kozak sequence at the N-terminus.

Forward Primer: 5′ TTAAAGCTTGCCACCATGGTCTTCACACTCGAA 3′
Reverse Primer: 3′ CAAGAATTCACTGTTGATGGTTACTCGGAACAG 3′

*Plasmid C*: The Lin28-LgBiT sequence was inserted into a pcDNA3 vector containing KpnI (fwd) and NotI (rev) restriction enzyme sites as previously described by our lab.[1] The Lin28 and LgBiT sequences were separated by an AsiSI restriction enzyme cut site. To clone C-terminal LgBiT constructs (RBP-LgBiT), the Lin28 sequence was removed, and the RBP-of-interest was inserted using appropriate restriction enzymes.

*LgBiT-MBNL1 cloning.* A synthetic human Muscleblind-like1 (MBNL1) gene fragment was purchased from Twist Bioscience and inserted into *Plasmid A* using standard cloning techniques with XhoI and XbaI restriction enzymes to produce an N-terminal LgBiT tag.

**Gene Fragment:**
5′CTCGAGATGGCAGTGTCCGTGACCCCGATAAGGGACACTAAGTGGCTTACT
CTGGAGGTGTGTCGAGAGTTCCAAAGAGGGACATGCTCTAGGCCCGACACTG
AATGCAAGTTTGCCCATCCCAGCAAAAGTTGCCAGGTCGAAAATGGACGTGT
TATAGCCTGCTTTGACAGCCTGAAGGGCCGTTGCTCACGGGAAAATTGTAAG
TATCTCCATCCACCCCCACACCTGAAGACACAGCTGGAGATCAACGGTCGCA
ACAACCTGATCCAGCAGAAGAACATGGCAATGCTGGCACAGCAGATGCAGC
TGGCCAATGCTATGATGCCTGGAGCCCCCCTCCAGCCCGTACCTATGTTCTCC
GTCGCCCCCTCTCTGGCGACAAACGCTTCCGCGGCCGCCTTTAATCCCTATCT
GGGACCTGTGTCCCCGTCACTCGTGCCAGCTGAGATTCTGCCGACCGCCCCA
ATGCTGGTGACCGGTAACCCCGGAGTCCCAGTCCCAGCGGCCGCAGCCGCAG
CTGCCCAGAAGCTGATGAGAACTGATAGGTTAGAGGTGTGCCGTGAGTATCA
ACGGGGCAATTGCAATAGGGGCGAGAACGACTGTCGGTTTGCGCACCCGGCA
GATTCCACCATGATAGATACCAATGACAACACAGTGACTGTATGCATGGACT
ACATTAAAGGACGTTGTTCAAGGGAAAAATGCAAGTATTTCCATCCTCCCGC
ACACCTCCAGGCGAAGATTAAGGCCGCCCAATACCAGGTTAACCAGGCGGCC
GCTGCTCAAGCAGCCGCCACAGCAGCAGCTATGGGTATTCCCCAGGCTGTGC
TGCCTCCGTTGCCTAAGAGACCTGCTTTGGAAAAGACTAACGGCGCAACAGC
CGTATTCAACACCGGAATCTTCCAATATCAGCAGGCACTGGCCAACATGCAG

CTGCAACAGCACACAGCTTTTCTGCCACCTGGGAGTATTCTTTGTATGACCCC
CGCGACGAGTGTCGTGCCCATGGTGCACGGAGCTACCCCCGCAACAGTCAGC
GCTGCTACAACGTCCGCCACTAGTGTGCCATTTGCCGCAACGGCTACCGCAA
ATCAAATACCTATCATCAGCGCTGAGCATCTGACTTCACATAAGTATGTAACC
CAGATGTAGTCTAGA3′

*MBNL1-LgBiT cloning*. A synthetic human MBNL1 gene fragment was purchased from Twist Bioscience and inserted into *Plasmid C* using standard cloning techniques with KpnI and AsiSI restriction enzymes to produce a C-terminal LgBiT tag. Gene fragment contains a Kozak sequence at the N-terminus.

**Gene Fragment:**
5′GGTACCGCCACCATGGCAGTGTCCGTGACCCCGATAAGGGACACTAAGTGG
CTTACTCTGGAGGTGTGTCGAGAGTTCCAAAGAGGGACATGCTCTAGGCCCG
ACACTGAATGCAAGTTTGCCCATCCCAGCAAAAGTTGCCAGGTCGAAAATGG
ACGTGTTATAGCCTGCTTTGACAGCCTGAAGGGCCGTTGCTCACGGGAAAAT
TGTAAGTATCTCCATCCACCCCCACACCTGAAGACACAGCTGGAGATCAACG
GTCGCAACAACCTGATCCAGCAGAAGAACATGGCAATGCTGGCACAGCAGA
TGCAGCTGGCCAATGCTATGATGCCTGGAGCCCCCCTCCAGCCCGTACCTATG
TTCTCCGTCGCCCCCTCTCTGGCGACAAACGCTTCCGCGGCCGCCTTTAATCC
CTATCTGGGACCTGTGTCCCCGTCACTCGTGCCAGCTGAGATTCTGCCGACCG
CCCCAATGCTGGTGACCGGTAACCCCGGAGTCCCAGTCCCAGCGGCCGCAGC
CGCAGCTGCCCAGAAGCTGATGAGAACTGATAGGTTAGAGGTGTGCCGTGAG
TATCAACGGGGCAATTGCAATAGGGGCGAGAACGACTGTCGGTTTGCGCACC
CGGCAGATTCCACCATGATAGATACCAATGACAACACAGTGACTGTATGCAT
GGACTACATTAAAGGACGTTGTTCAAGGGAAAAATGCAAGTATTTCCATCCT
CCCGCACACCTCCAGGCGAAGATTAAGGCCGCCCAATACCAGGTTAACCAGG
CGGCCGCTGCTCAAGCAGCCGCCACAGCAGCAGCTATGGGTATTCCCCAGGC
TGTGCTGCCTCCGTTGCCTAAGAGACCTGCTTTGGAAAAGACTAACGGCGCA
ACAGCCGTATTCAACACCGGAATCTTCCAATATCAGCAGGCACTGGCCAACA
TGCAGCTGCAACAGCACACAGCTTTTCTGCCACCTGGGAGTATTCTTTGTATG
ACCCCCGCGACGAGTGTCGTGCCCATGGTGCACGGAGCTACCCCCGCAACAG
TCAGCGCTGCTACAACGTCCGCCACTAGTGTGCCATTTGCCGCAACGGCTACC
GCAAATCAAATACCTATCATCAGCGCTGAGCATCTGACTTCACATAAGTATGT
AACCCAGATGGCGATCGC3′

*MBNL1-260-LgBiT cloning*. A truncated MBNL1 containing the first 260 amino acids from the human gene sequence was PCR amplified using MBNL1-LgBiT as a template. MBNL1-260 was inserted into *Plasmid C* using standard cloning techniques with KpnI and AsiSI restriction enzymes to produce a C-terminal LgBiT tag. Primer inserts Kozak sequence at the N-terminus.

Forward Primer: 5′ TTAGGTACCGCCACCATGGCAGTGTCCGTGACC 3′
Reverse Primer: 5′ TAAGCGATCGCAGCAGCGGCCGCCTGG 3′

**Truncated Sequence**:

5′GGTACCGCCACCATGGCAGTGTCCGTGACCCCGATAAGGGACACTAAGTGG
CTTACTCTGGAGGTGTGTCGAGAGTTCCAAAGAGGGACATGCTCTAGGCCCG
ACACTGAATGCAAGTTTGCCCATCCCAGCAAAAGTTGCCAGGTCGAAAATGG
ACGTGTTATAGCCTGCTTTGACAGCCTGAAGGGCCGTTGCTCACGGAAAAT
TGTAAGTATCTCCATCCACCCCCACACCTGAAGACACAGCTGGAGATCAACG
GTCGCAACAACCTGATCCAGCAGAAGAACATGGCAATGCTGGCACAGCAGA
TGCAGCTGGCCAATGCTATGATGCCTGGAGCCCCCCTCCAGCCCGTACCTATG
TTCTCCGTCGCCCCCTCTCTGGCGACAAACGCTTCCGCGGCCGCCTTTAATCC
CTATCTGGGACCTGTGTCCCCGTCACTCGTGCCAGCTGAGATTCTGCCGACCG
CCCCAATGCTGGTGACCGGTAACCCCGGAGTCCCAGTCCCAGCGGCCGCAGC
CGCAGCTGCCCAGAAGCTGATGAGAACTGATAGGTTAGAGGTGTGCCGTGAG
TATCAACGGGGCAATTGCAATAGGGGCGAGAACGACTGTCGGTTTGCGCACC
CGGCAGATTCCACCATGATAGATACCAATGACAACACAGTGACTGTATGCAT
GGACTACATTAAAGGACGTTGTTCAAGGGAAAAATGCAAGTATTTCCATCCT
CCCGCACACCTCCAGGCGAAGATTAAGGCCGCCCAATACCAGGTTAACCAGG
CGGCCGCTGCTGCGATCGC3′

*MBNL1-ZnF12-LgBiT cloning*. A truncated MBNL1 containing the Zinc Finger Domains 1 and 2 from the human gene sequence was PCR amplified using MBNL1-LgBiT as a template. MBNL1-ZnF12 was inserted into Plasmid C using standard cloning techniques with KpnI and AsiSI restriction enzymes to produce a C-terminal LgBiT tag. Forward primer inserts Kozak sequence at the N-terminus.

Forward Primer: 5′ TTAGGTACCGCCACCATGGCAGTGTCCGTGACC 3′
Reverse Primer: 5′ TGAGCGATCGCCTGCTGGATCAGGTTGTT 3′

**Truncated Sequence**:
5′GGTACCGCCACCATGGCAGTGTCCGTGACCCCGATAAGGGACACTAAGTGG
CTTACTCTGGAGGTGTGTCGAGAGTTCCAAAGAGGGACATGCTCTAGGCCCG
ACACTGAATGCAAGTTTGCCCATCCCAGCAAAAGTTGCCAGGTCGAAAATGG
ACGTGTTATAGCCTGCTTTGACAGCCTGAAGGGCCGTTGCTCACGGAAAAT
TGTAAGTATCTCCATCCACCCCCACACCTGAAGACACAGCTGGAGATCAACG
GTCGCAACAACCTGATCCAGCAGGCGATCGC3′

*MBNL1-ZnF34-LgBiT cloning*. A truncated MBNL1 containing the Zinc Finger Domains 3 and 4 from the human gene sequence was PCR amplified using MBNL1-LgBiT as a template. MBNL1-ZnF34 was inserted into Plasmid C using standard cloning techniques with KpnI and AsiSI restriction enzymes to produce a C-terminal LgBiT tag. Forward primer inserts a Kozak sequence at the N-terminus.

Forward Primer: 5′ TCAGGTACCGCCACCATG AGAACTGATAGGTTAGAG 3′
Reverse Primer: 5′ TGAGCGATCGCCTGCTGGATCAGGTTGTT 3′

**Truncated Sequence:**
5′GGTACCGCCACCATGAGAACTGATAGGTTAGAGGTGTGCCGTGAGTATCAA
CGGGGCAATTGCAATAGGGGCGAGAACGACTGTCGGTTTGCGCACCCGGCAG

ATTCCACCATGATAGATACCAATGACAACACAGTGACTGTATGCATGGACTA
CATTAAAGGACGTTGTTCAAGGGAAAAATGCAAGTATTTCCATCCTCCCGCA
CACCTCCAGGCGAAGATTAAGGCCGCCCAATACCAGGTTAACCAGGCGGCCG
CTGCTGCGATCGC3′

*LgBiT-Stau1 cloning*. A synthetic human Staufen1 (Stau1) gene fragment was PCR amplified from HeLa cDNA made from random hexamers and inserted into *Plasmid A* using standard cloning techniques with XhoI and XbaI restriction enzymes to produce an N-terminal LgBiT tag.

Forward Primer: 5′ ACTACTCGAGATGTCTCAAGTTCAAGTG 3′
Reverse Primer: 5′ CCTCTAGATCAGCACCTCCCACACAC 3′

**Gene Sequence**:
5′CTCGAGATGTCTCAAGTTCAAGTGCAAGTTCAGAACCCATCTGCTGCTCTCT
CAGGGAGCCAAATACTGAACAAGAACCAGTCTCTTCTCTCACAGCCTTTGAT
GAGTATTCCTTCTACTACTAGCTCTCTGCCCTCTGAAAATGCAGGTAGACCCA
TTCAAAACTCTGCTTTACCCTCTGCATCTATTACATCCACCAGTGCAGCTGCA
GAAAGCATAACCCCTACTGTAGAACTAAATGCACTGTGCATGAAACTTGGAA
AAAAACCAATGTATAAGCCTGTTGACCCTTACTCTCGGATGCAGTCCACCTAT
AACTACAACATGAGAGGAGGTGCTTATCCCCCGAGGTACTTTTACCCATTTCC
AGTTCCACCTTTACTTTATCAAGTGGAACTTTCTGTGGGAGGACAGCAATTTA
ATGGCAAAGGAAAGACAAGACAGGCTGCGAAACACGATGCTGCTGCCAAAG
CGTTGAGGATCCTGCAGAATGAGCCCCTGCCAGAGAGGCTGGAGGTGAATGG
AAGAGAATCCGAAGAAGAAAATCTCAATAAATCTGAAATAAGTCAAGTGTTT
GAGATTGCACTTAAACGGAACTTGCCTGTGAATTTCGAGGTGGCCCGGGAGA
GTGGCCCACCCCACATGAAGAACTTTGTGACCAAGGTTTCGGTTGGGGAGTT
TGTGGGGGAAGGTGAAGGGAAAAGCAAGAAGATTTCAAAGAAAAATGCCGC
CATAGCTGTTCTTGAGGAGCTGAAGAAGTTACCGCCCCTGCCTGCAGTTGAA
CGAGTAAAGCCTAGAATCAAAAAGAAAACAAAACCCATAGTCAAGCCACAG
ACAAGCCCAGAATATGGCCAGGGGATCAATCCGATTAGCCGACTGGCCCAGA
TCCAGCAGGCAAAAAAGGAGAAGGAGCCAGAGTACACGCTCCTCACAGAGC
GAGGCCTCCCGCGCCGCAGGGAGTTTGTGATGCAGGTGAAGGTTGGAAACCA
CACTGCAGAAGGAACGGGCACCAACAAGAAGGTGGCCAAGCGCAATGCAGC
CGAGAACATGCTGGAGATCCTTGGTTTCAAAGTCCCGCAGGCGCAGCCCACC
AAACCCGCACTCAAGTCAGAGGAGAAGACACCCATAAAGAAACCAGGGGAT
GGAAGAAAAGTAACCTTTTTTGAACCTGGCTCTGGGGATGAAAATGGGACTA
GTAATAAAGAGGATGAGTTCAGGATGCCTTATCTAAGTCATCAGCAGCTGCC
TGCTGGAATTCTTCCCATGGTGCCCGAGGTCGCCCAGGCTGTAGGAGTTAGTC
AAGGACATCACACCAAAGATTTTACCAGGGCAGCTCCGAATCCTGCCAAGGC
CACGGTAACTGCCATGATAGCCCGAGAGTTGTTGTATGGGGGCACCTCGCCC
ACAGCCGAGACCATTTTAAAGAATAACATCTCTTCAGGCCACGTACCCCATG
GACCTCTCACGAGACCCTCTGAGCAACTGGACTATCTTTCCAGAGTCCAGGG
ATTCCAGGTTGAATACAAAGACTTCCCCAAAAACAACAAGAACGAATTTGTA
TCTCTTATCAATTGCTCCTCTCAGCCACCTCTGATCAGCCATGGTATCGGCAA
GGATGTGGAGTCCTGCCATGATATGGCTGCGCTGAACATCTTAAAGTTGCTGT

CTGAGTTGGACCAACAAAGTACAGAGATGCCAAGAACAGGAAACGGACCAA
TGTCTGTGTGTGGGAGGTGCTGATCTAGA3′

*Stau1-LgBiT cloning*. A synthetic human Staufen1 (Stau1) gene fragment was PCR amplified from LgBiT-Stau1 and inserted into *Plasmid C* using standard cloning techniques with KpnI and AsiSI restriction enzymes to produce a C-terminal LgBiT tag. Forward primer inserts a Kozak sequence at the N-terminus.

Forward Primer: 5′ ATGGTACCGCCACCATGTCTCAAGTTCAAGTG 3′
Reverse Primer: 5′ ATGCGATCGCGCACCTCCCACACACAGA 3′

**Gene Sequence**:
5′GGTACCGCCACCATGTCTCAAGTTCAAGTGCAAGTTCAGAACCCATCTGCT
GCTCTCTCAGGGAGCCAAATACTGAACAAGAACCAGTCTCTTCTCTCACAGC
CTTTGATGAGTATTCCTTCTACTACTAGCTCTCTGCCCTCTGAAAATGCAGGT
AGACCCATTCAAAACTCTGCTTTACCCTCTGCATCTATTACATCCACCAGTGC
AGCTGCAGAAAGCATAACCCCTACTGTAGAACTAAATGCACTGTGCATGAAA
CTTGGAAAAAAACCAATGTATAAGCCTGTTGACCCTTACTCTCGGATGCAGTC
CACCTATAACTACAACATGAGAGGAGGTGCTTATCCCCCGAGGTACTTTTACC
CATTTCCAGTTCCACCTTTACTTTATCAAGTGGAACTTTCTGTGGGAGGACAG
CAATTTAATGGCAAAGGAAAGACAAGACAGGCTGCGAAACACGATGCTGCT
GCCAAAGCGTTGAGGATCCTGCAGAATGAGCCCCTGCCAGAGAGGCTGGAG
GTGAATGGAAGAGAATCCGAAGAAGAAAATCTCAATAAATCTGAAATAAGT
CAAGTGTTTGAGATTGCACTTAAACGGAACTTGCCTGTGAATTTCGAGGTGGC
CCGGGAGAGTGGCCCACCCCACATGAAGAACTTTGTGACCAAGGTTTCGGTT
GGGGAGTTTGTGGGGGAAGGTGAAGGGAAAAGCAAGAAGATTTCAAAGAAA
AATGCCGCCATAGCTGTTCTTGAGGAGCTGAAGAAGTTACCGCCCCTGCCTG
CAGTTGAACGAGTAAAGCCTAGAATCAAAAAGAAAACAAAACCCATAGTCA
AGCCACAGACAAGCCCAGAATATGGCCAGGGGATCAATCCGATTAGCCGACT
GGCCCAGATCCAGCAGGCAAAAAAGGAGAAGGAGCCAGAGTACACGCTCCT
CACAGAGCGAGGCCTCCCGCGCCGCAGGGAGTTTGTGATGCAGGTGAAGGTT
GGAAACCACACTGCAGAAGGAACGGGCACCAACAAGAAGGTGGCCAAGCGC
AATGCAGCCGAGAACATGCTGGAGATCCTTGGTTTCAAAGTCCCGCAGGCGC
AGCCCACCAAACCCGCACTCAAGTCAGAGGAGAAGACACCCATAAAGAAC
CAGGGGATGGAAGAAAAGTAACCTTTTTGAACCTGGCTCTGGGGATGAAAA
TGGGACTAGTAATAAAGAGGATGAGTTCAGGATGCCTTATCTAAGTCATCAG
CAGCTGCCTGCTGGAATTCTTCCCATGGTGCCCGAGGTCGCCCAGGCTGTAG
GAGTTAGTCAAGGACATCACACCAAAGATTTTACCAGGGCAGCTCCGAATCC
TGCCAAGGCCACGGTAACTGCCATGATAGCCCGAGAGTTGTTGTATGGGGGC
ACCTCGCCCACAGCCGAGACCATTTTAAAGAATAACATCTCTTCAGGCCACG
TACCCCATGGACCTCTCACGAGACCCTCTGAGCAACTGGACTATCTTTCCAGA
GTCCAGGGATTCCAGGTTGAATACAAAGACTTCCCCAAAAACAACAAGAACG
AATTTGTATCTCTTATCAATTGCTCCTCTCAGCCACCTCTGATCAGCCATGGTA
TCGGCAAGGATGTGGAGTCCTGCCATGATATGGCTGCGCTGAACATCTTAAA
GTTGCTGTCTGAGTTGGACCAACAAAGTACAGAGATGCCAAGAACAGGAAAC
GGACCAATGTCTGTGTGTGGGAGGTGCGCGATCGC3′

*LgBiT-CELF1 cloning*. A synthetic human CUGBP ELAV-like factor 1 (CELF1) gene fragment was purchased from Twist Bioscience and inserted into *Plasmid A* using standard cloning techniques with XhoI and XbaI to produce an N-terminal LgBiT tag.

**Gene Sequence**:
5′CTCGAGATGAACGGCACCCTGGACCACCCAGACCAACCAGATCTTGATGCT
ATCAAGATGTTTGTGGGCCAGGTTCCAAGGACCTGGTCTGAAAAGGACTTGC
GGGAACTCTTCGAACAGTATGGTGCTGTGTATGAAATCAACGTCCTAAGGGA
TAGGAGCCAAAACCCGCCTCAGAGCAAAGGGTGCTGTTTTGTTACATTTTAC
ACCCGTAAAGCTGCATTAGAAGCTCAGAATGCTCTTCACAACATGAAAGTCC
TCCCAGGGATGCATCACCCTATACAGATGAAACCTGCTGACAGTGAGAAGAA
CAATGCAGTGGAAGACAGGAAGCTGTTTATTGGTATGATTTCCAAGAAGTGC
ACTGAAAATGACATCCGAGTCATGTTCTCTTCGTTTGGACAGATTGAAGAATG
CCGGATATTGCGGGGACCTGATGGCCTGAGCCGAGGTTGTGCATTTGTGACTT
TTACAACAAGAGCCATGGCACAGACGGCTATCAAGGCAATGCACCAAGCAC
AGACCATGGAGGGTTGCTCATCACCCATGGTGGTAAAATTTGCTGATACACA
GAAGGACAAAGAACAGAAGAGAATGGCCCAGCAGCTCCAGCAGCAGATGCA
GCAAATCAGCGCAGCATCTGTGTGGGGAAACCTTGCTGGTCTAAATACTCTT
GGACCCCAGTATTTAGCACTCCTTCAGCAGACTGCCTCCTCTGGGAACCTCAA
CACCCTGAGCAGCCTCCACCCAATGGGAGGGTTGAATGCAATGCAGTTACAG
AATTTGGCTGCACTAGCTGCTGCAGCTAGTGCAGCTCAGAACACACCAAGTG
GTACCAATGCTCTCACTACATCCAGCAGTCCCCTCAGCGTGCTCACTAGTTCA
GGGTCCTCACCTAGCTCTAGCAGCAGTAATTCTGTCAACCCCATAGCCTCACT
TGGAGCCCTGCAGACATTAGCTGGAGCAACGGCTGGCCTCAATGTTGGCTCT
TTGGCAGGAATGGCTGCTTTAAATGGTGGCCTGGGCAGCAGTGGCCTTTCCA
ATGGCACCGGGAGCACCATGGAGGCCCTCACTCAGGCCTACTCGGGTATCCA
GCAATATGCTGCTGCTGCGCTCCCCACTCTGTACAACCAGAATCTTCTGACAC
AGCAGAGTATTGGTGCTGCTGGAAGCCAGAAGGAAGGTCCAGAGGGAGCCA
ACCTGTTCATCTACCACCTGCCCCAGGAGTTTGGTGATCAGGACCTGCTGCAG
ATGTTTATGCCCTTTGGGAATGTCGTGTCTGCCAAGGTTTTCATAGACAAGCA
GACAAACCTGAGCAAGTGTTTTGGTTTTGTAAGTTACGACAATCCTGTTTCGG
CCCAAGCTGCCATCCAGTCCATGAACGGCTTTCAGATTGGCATGAAGCGGCT
TAAAGTGCAGCTCAAACGTTCGAAGAATGACAGCAAGCCCTACTGATCTAGA
3′

*CELF1-LgBiT cloning*: A synthetic human CELF1 gene fragment was purchased from Twist Bioscience and inserted into *Plasmid C* using standard cloning techniques with KpnI and AsiSI restriction enzymes to produce a C-terminal LgBiT tag. Gene fragment contains a Kozak sequence at the N-terminus.

**Gene Sequence**:
5′GGTACCGCCACCATGAACGGCACCCTGGACCACCCAGACCAACCAGATCTT
GATGCTATCAAGATGTTTGTGGGCCAGGTTCCAAGGACCTGGTCTGAAAAGG
ACTTGCGGGAACTCTTCGAACAGTATGGTGCTGTGTATGAAATCAACGTCCTA
AGGGATAGGAGCCAAAACCCGCCTCAGAGCAAAGGGTGCTGTTTTGTTACAT

TTTACACCCGTAAAGCTGCATTAGAAGCTCAGAATGCTCTTCACAACATGAA
AGTCCTCCCAGGGATGCATCACCCTATACAGATGAAACCTGCTGACAGTGAG
AAGAACAATGCAGTGGAAGACAGGAAGCTGTTTATTGGTATGATTTCCAAGA
AGTGCACTGAAAATGACATCCGAGTCATGTTCTCTTCGTTTGGACAGATTGAA
GAATGCCGGATATTGCGGGGACCTGATGGCCTGAGCCGAGGTTGTGCATTTG
TGACTTTTACAACAAGAGCCATGGCACAGACGGCTATCAAGGCAATGCACCA
AGCACAGACCATGGAGGGTTGCTCATCACCCATGGTGGTAAAATTTGCTGAT
ACACAGAAGGACAAAGAACAGAAGAGAATGGCCCAGCAGCTCCAGCAGCAG
ATGCAGCAAATCAGCGCAGCATCTGTGTGGGGAAACCTTGCTGGTCTAAATA
CTCTTGGACCCCAGTATTTAGCACTCCTTCAGCAGACTGCCTCCTCTGGGAAC
CTCAACACCCTGAGCAGCCTCCACCCAATGGGAGGGTTGAATGCAATGCAGT
TACAGAATTTGGCTGCACTAGCTGCTGCAGCTAGTGCAGCTCAGAACACACC
AAGTGGTACCAATGCTCTCACTACATCCAGCAGTCCCCTCAGCGTGCTCACTA
GTTCAGGGTCCTCACCTAGCTCTAGCAGCAGTAATTCTGTCAACCCCATAGCC
TCACTTGGAGCCCTGCAGACATTAGCTGGAGCAACGGCTGGCCTCAATGTTG
GCTCTTTGGCAGGAATGGCTGCTTTAAATGGTGGCCTGGGCAGCAGTGGCCTT
TCCAATGGCACCGGGAGCACCATGGAGGCCCTCACTCAGGCCTACTCGGGTA
TCCAGCAATATGCTGCTGCTGCGCTCCCCACTCTGTACAACCAGAATCTTCTG
ACACAGCAGAGTATTGGTGCTGCTGGAAGCCAGAAGGAAGGTCCAGAGGGA
GCCAACCTGTTCATCTACCACCTGCCCCAGGAGTTTGGTGATCAGGACCTGCT
GCAGATGTTTATGCCCTTTGGGAATGTCGTGTCTGCCAAGGTTTTCATAGACA
AGCAGACAAACCTGAGCAAGTGTTTTGGTTTTGTAAGTTACGACAATCCTGTT
TCGGCCCAAGCTGCCATCCAGTCCATGAACGGCTTTCAGATTGGCATGAAGC
GGCTTAAAGTGCAGCTCAAACGTTCGAAGAATGACAGCAAGCCCTACGCGAT
CGC3′

*LgBiT-Msi1 cloning*. A synthetic human Musashi1 (Msi1) gene fragment was purchased from Twist Bioscience and inserted into *Plasmid B* using standard cloning techniques with EcoRV and XbaI restriction enzymes to produce an N-terminal LgBiT tag.

**Gene Fragment**:
5′GATATCTTATGGAGACTGACGCGCCCCAGCCCGGCCTCGCCTCCCCGGACT
CGCCGCACGACCCCTGCAAGATGTTCATCGGGGGACTCAGTTGGCAGACTAC
GCAGGAAGGGCTGCGCGAATACTTCGGCCAGTTCGGGGAGGTGAAGGAGTG
TCTGGTGATGCGGGACCCCCTGACCAAGAGATCCAGGGGTTTCGGCTTCGTC
ACTTTCATGGACCAGGCGGGGGTGGATAAAGTGCTGGCGCAATCGCGGCACG
AGCTCGACTCCAAAACAATTGACCCTAAGGTGGCCTTCCCTCGGCGAGCACA
GCCCAAGATGGTGACTCGAACGAAGAAGATCTTTGTGGGGGGGCTGTCGGTG
AACACCACGGTGGAGGACGTGAAGCAATATTTTGAGCAGTTTGGGAAGGTGG
ACGACGCCATGCTGATGTTTGACAAAACCACCAACCGGCACCGAGGGTTCGG
GTTTGTCACGTTTGAGAGTGAGGACATCGTGGAGAAAGTGTGTGAAATTCAT
TTTCATGAAATCAACAACAAAATGGTGGAATGTAAGAAAGCTCAGCCAAAGG
AGGTGATGTCGCCAACGGGCTCAGCCCGGGGGAGGTCTCGAGTCATGCCCTA
CGGAATGGACGCCTTCATGCTGGGCATCGGCATGCTGGGTTACCCAGGTTTCC
AAGCCACAACCTACGCCAGCCGGAGTTATACAGGCCTCGCCCCTGGCTACAC
CTACCAGTTCCCCGAATTCCGTGTAGAGCGGACCCCTCTCCCGAGCGCCCCA

GTCCTCCCCGAGCTTACAGCCATTCCTCTCACTGCCTACGGACCAATGGCGGC
GGCAGCGGCGGCAGCGGCTGTGGTTCGAGGGACAGGCTCTCACCCCTGGACG
ATGGCTCCCCCTCCAGGTTCGACTCCCAGCCGCACAGGGGGCTTCCTGGGGA
CCACCAGCCCCGGCCCCATGGCCGAGCTCTACGGGGCGGCCAACCAGGACTC
GGGGGTCAGCAGTTACATCAGCGCCGCCAGCCCTGCCCCCAGCACCGGCTTC
GGCCACAGTCTTGGGGGCCCTTTGATTGCCACAGCCTTCACCAATGGGTACC
ACTGATCTAGA3′

*Msi1-LgBiT cloning*. A synthetic human Musashi1 (Msi1) gene fragment was purchased from Twist Bioscience and inserted into *Plasmid C* using standard cloning techniques with HindIII and AsiSI restriction enzymes to produce a C-terminal LgBiT tag. Gene fragment contains a Kozak sequence at the N-terminus.

**Gene Fragment**:
5′AAGCTTGCCACCATGGAGACTGACGCGCCCCAGCCCGGCCTCGCCTCCCCG
GACTCGCCGCACGACCCCTGCAAGATGTTCATCGGGGGACTCAGTTGGCAGA
CTACGCAGGAAGGGCTGCGCGAATACTTCGGCCAGTTCGGGGAGGTGAAGG
AGTGTCTGGTGATGCGGGACCCCCTGACCAAGAGATCCAGGGGTTTCGGCTT
CGTCACTTTCATGGACCAGGCGGGGGTGGATAAAGTGCTGGCGCAATCGCGG
CACGAGCTCGACTCCAAAACAATTGACCCTAAGGTGGCCTTCCCTCGGCGAG
CACAGCCCAAGATGGTGACTCGAACGAAGAAGATCTTTGTGGGGGGGCTGTC
GGTGAACACCACGGTGGAGGACGTGAAGCAATATTTTGAGCAGTTTGGGAAG
GTGGACGACGCCATGCTGATGTTTGACAAAACCACCAACCGGCACCGAGGGT
TCGGGTTTGTCACGTTTGAGAGTGAGGACATCGTGGAGAAAGTGTGTGAAAT
TCATTTTCATGAAATCAACAACAAAATGGTGGAATGTAAGAAAGCTCAGCCA
AAGGAGGTGATGTCGCCAACGGGCTCAGCCCGGGGGAGGTCTCGAGTCATGC
CCTACGGAATGGACGCCTTCATGCTGGGCATCGGCATGCTGGGTTACCCAGG
TTTCCAAGCCACAACCTACGCCAGCCGGAGTTATACAGGCCTCGCCCCTGGC
TACACCTACCAGTTCCCCGAATTCCGTGTAGAGCGGACCCCTCTCCCGAGCGC
CCCAGTCCTCCCCGAGCTTACAGCCATTCCTCTCACTGCCTACGGACCAATGG
CGGCGGCAGCGGCGGCAGCGGCTGTGGTTCGAGGGACAGGCTCTCACCCCTG
GACGATGGCTCCCCCTCCAGGTTCGACTCCCAGCCGCACAGGGGGCTTCCTG
GGGACCACCAGCCCCGGCCCCATGGCCGAGCTCTACGGGGCGGCCAACCAG
GACTCGGGGGTCAGCAGTTACATCAGCGCCGCCAGCCCTGCCCCCAGCACCG
GCTTCGGCCACAGTCTTGGGGGCCCTTTGATTGCCACAGCCTTCACCAATGGG
TACCACGCGATCGC3′

*LgBiT-Msi2 cloning*. Human Musashi2 (Msi2) was PCR amplified from a purchased plasmid, Msi2 variant 1 in pFN21A (Promega), and inserted into *Plasmid A* using standard cloning techniques with XhoI and XbaI to produce an N-terminal LgBiT tag.

Forward Primer: 5′ TCTCCTCGAGATGGAGGCAAATGGGAGCC 3′
Reverse Primer: 5′ CAGTGAATTCTCAATGGTATCCATTTGTAAAGGCCG 3′

**Gene Sequence**:

5′CTCGAGATGGAGGCAAATGGAGCCAAGGCACCTCGGGCAGCGCCAACGA
CTCCCAGCACGACCCCGGTAAAATGTTTATCGGTGGACTGAGCTGGCAGACC
TCACCAGATAGCCTTAGAGACTATTTTAGCAAATTTGGAGAAATTAGAGAAT
GTATGGTCATGAGAGATCCCACTACGAAACGCTCCAGAGGCTTCGGTTTCGT
CACGTTCGCAGACCCAGCAAGTGTAGATAAAGTATTAGGTCAGCCCCACCAT
GAGTTAGATTCCAAGACGATTGACCCCAAAGTTGCATTTCCTCGTCGAGCGC
AACCCAAGATGGTCACAAGAACAAAGAAAATATTTGTAGGCGGGTTATCTGC
GAACACAGTAGTGGAAGATGTAAAGCAATATTTCGAGCAGTTTGGCAAGGTG
GAAGATGCAATGCTGATGTTTGATAAAACTACCAACAGGCACAGAGGGTTTG
GCTTTGTCACTTTTGAGAATGAAGATGTTGTGGAGAAAGTCTGTGAGATTCAT
TTCCATGAAATCAATAATAAAATGGTAGAATGTAAGAAAGCTCAGCCGAAAG
AAGTCATGTTCCCACCTGGGACAAGAGGCCGGGCCCGGGGACTGCCTTACAC
CATGGACGCGTTCATGCTTGGCATGGGGATGCTGGGATATCCCAACTTCGTG
GCGACCTATGGCCGTGGCTACCCCGGATTTGCTCCAAGCTATGGCTATCAGTT
CCCAGGCTTCCCAGCAGCGGCTTATGGACCAGTGGCAGCAGCGGCGGTGGCG
GCAGCAAGAGGATCAGGCTCCAACCCGGCGCGGCCCGGAGGCTTCCCGGGG
GCCAACAGCCCAGGACCTGTCGCCGATCTCTACGGCCCTGCCAGCCAGGACT
CCGGAGTGGGGAATTACATAAGTGCGGCTAGCCCACAGCCGGGCTCGGGCTT
CGGCCACGGCATAGCTGGACCTTTGATTGCAACGGCCTTTACAAATGGATAC
CATTGAGAATTCTGCAGATATCCATCACACTGGCGGCCGCTCGAGCATGCAT
CTAGA3′

*Msi2-LgBiT cloning*. Human Msi2 was PCR amplified from a purchased plasmid, Msi2 variant 1 in pFN21A (Promega), and inserted into *Plasmid C* using standard cloning techniques with KpnI and AsiSI to produce a C-terminal LgBiT tag. Forward primer inserts a Kozak sequence at the N-terminus.

Forward Primer: 5′ GTACGGTACCGCCACCATGGAGGCAAATGGAGCCAAG 3′
Reverse Primer: 5′ GTCACGGCGATCGCATGGTATCCATTTGTAAAGGCC 3′

**Gene Fragment**:
5′GGTACCGCCACCATGGAGGCAAATGGAGCCAAGGCACCTCGGGCAGCGC
CAACGACTCCCAGCACGACCCCGGTAAAATGTTTATCGGTGGACTGAGCTGG
CAGACCTCACCAGATAGCCTTAGAGACTATTTTAGCAAATTTGGAGAAATTA
GAGAATGTATGGTCATGAGAGATCCCACTACGAAACGCTCCAGAGGCTTCGG
TTTCGTCACGTTCGCAGACCCAGCAAGTGTAGATAAAGTATTAGGTCAGCCC
CACCATGAGTTAGATTCCAAGACGATTGACCCCAAAGTTGCATTTCCTCGTCG
AGCGCAACCCAAGATGGTCACAAGAACAAAGAAAATATTTGTAGGCGGGTT
ATCTGCGAACACAGTAGTGGAAGATGTAAAGCAATATTTCGAGCAGTTTGGC
AAGGTGGAAGATGCAATGCTGATGTTTGATAAAACTACCAACAGGCACAGAG
GGTTTGGCTTTGTCACTTTTGAGAATGAAGATGTTGTGGAGAAAGTCTGTGAG
ATTCATTTCCATGAAATCAATAATAAAATGGTAGAATGTAAGAAAGCTCAGC
CGAAAGAAGTCATGTTCCCACCTGGGACAAGAGGCCGGGCCCGGGGACTGC
CTTACACCATGGACGCGTTCATGCTTGGCATGGGGATGCTGGGATATCCCAA
CTTCGTGGCGACCTATGGCCGTGGCTACCCCGGATTTGCTCCAAGCTATGGCT
ATCAGTTCCCAGGCTTCCCAGCAGCGGCTTATGGACCAGTGGCAGCAGCGGC

GGTGGCGGCAGCAAGAGGATCAGGCTCCAACCCGGCGCGGCCCGGAGGCTT
CCCGGGGGCCAACAGCCCAGGACCTGTCGCCGATCTCTACGGCCCTGCCAGC
CAGGACTCCGGAGTGGGGAATTACATAAGTGCGGCCAGCCCACAGCCGGGCT
CGGGCTTCGGCCACGGCATAGCTGGACCTTTGATTGCAACGGCCTTTACAAAT
GGATACCATGCGATCGC3′

*LgBiT-AUF1 P37 cloning*. A synthetic human heterogenous ribonucleoprotein D (hnRNP D; also known as AU-binding Factor 1, AUF1) P37 gene fragment was purchased from Twist Bioscience and inserted into *Plasmid A* using standard cloning techniques with XhoI and ApaI restriction enzymes to produce an N-terminal LgBiT tag.

**Gene Fragment**:
5′CTCGAGATGTCTGAAGAACAATTTGGTGGTGATGGTGCCGCTGCAGCCGCC
ACAGCAGCTGTTGGCGGTTCAGCAGGTGAACAAGAAGGCGCAATGGTTGCAG
CTACTCAAGGAGCCGCAGCCGCAGCAGGCTCTGGCGCTGGAACGGGAGGAG
GTACGGCTTCCGGTGGAACAGAGGGTGGATCCGCGGAATCCGAAGGAGCCA
AAATCGATGCAAGCAAGAATGAAGAAGACGAGGGTAAGATGTTCATTGGCG
GTCTGTCATGGGATACGACCAAGAAGGACCTCAAAGATTATTTCAGTAAGTT
CGGCGAGGTGGTGGATTGTACCCTTAAACTTGACCCAATTACCGGCAGGTCT
CGCGGGTTCGGGTTCGTTCTGTTCAAGGAGAGTGAATCTGTCGACAAAGTTAT
GGACCAGAAGGAGCACAAGCTCAACGGCAAAGTAATAGACCCGAAGAGAGC
AAAGGCTATGAAGACCAAGGAACCTGTGAAGAAGATCTTCGTGGGCGGTCTG
AGCCCTGACACCCCAGAGGAAAAGATTAGAGAATATTTCGGCGGCTTCGGCG
AAGTAGAGTCAATTGAATTACCAATGGATAATAAAACAAACAAACGGCGAG
GATTTTGTTTCATAACATTCAAAGAGGAAGAGCCGGTCAAGAAAATCATGGA
GAAGAAGTATCATAACGTCGGGCTGTCCAAGTGCGAGATTAAGGTTGCTATG
TCTAAAGAGCAGTACCAACAGCAACAGCAATGGGGATCACGCGGCGGGTTC
GCCGGCCGTGCTAGAGGGCGCGGCGGAGATCAACAATCAGGATACGGAAAA
GTGTCTAGACGGGGCGGCCACCAGAACTCCTATAAGCCTTATTAGGGGCCC3′

*AUF1-LgBiT P37 cloning*. A synthetic human AUF1 P37 gene fragment was purchased from Twist Bioscience and inserted into *Plasmid C* using standard cloning techniques with KpnI and AsiSI restriction enzymes to produce a C-terminal LgBiT tag. Gene fragment contains a Kozak sequence at the N-terminus.

**Gene Fragment**:
5′GGTACCGCCACCATGAGCGAAGAACAATTTGGAGGAGATGGAGCAGCCGC
TGCTGCTACAGCCGCCGTCGGTGGGAGTGCAGGTGAACAAGAAGGCGCAATG
GTTGCTGCCACTCAAGGAGCTGCAGCTGCCGCCGGGTCAGGCGCTGGTACAG
GAGGAGGAACAGCCAGCGGTGGAACTGAGGGCGGATCAGCTGAATCCGAAG
GTGCCAAAATCGATGCTTCTAAGAATGAAGAAGACGAGGGCAAGATGTTCAT
CGGCGGACTGTCTTGGGATACAACTAAGAAGGACCTTAAAGATTATTTCAGC
AAGTTCGGCGAGGTGGTCGATTGTACCTTAAAACTGGACCCCATTACCGGCC
GCTCCCGGGGCTTCGGGTTCGTCCTTTTCAAGGAGTCCGAATCCGTCGACAAA
GTTATGGACCAGAAGGAGCACAAGCTGAACGGAAAAGTCATCGACCCAAAG
CGGGCAAAGGCTATGAAGACGAAGGAACCCGTAAAGAAGATCTTCGTAGGC

GGTCTGAGTCCTGACACTCCCGAGGAAAGATCCGCGAATATTTCGGCGGGT
TCGGCGAAGTAGAGTCAATTGAACTGCCGATGGATAATAAAACTAACAAACG
GCGCGGCTTTTGTTTCATCACTTTCAAAGAAGAGGAGCCCGTCAAGAAAATC
ATGGAGAAGAAGTATCATAACGTGGGGTTGTCTAAGTGCGAGATCAAGGTCG
CTATGAGCAAAGAGCAGTACCAACAGCAACAGCAATGGGGCTCACGGGGCG
GGTTCGCCGGGCGGGCCAGAGGCCGGGGCGGCGATCAACAAAGCGGCTACG
GAAAAGTGAGTAGACGGGGTGGCCACCAGAACTCTTATAAGCCTTATGCGAT
CGC3′

*LgBiT-AUF1 P40 cloning*. A synthetic human AUF1 P40 gene fragment was purchased from Twist Bioscience and inserted into *Plasmid A* using standard cloning techniques with XhoI and ApaI restriction enzymes to produce an N-terminal LgBiT tag.

**Gene Fragment**:
5′CTCGAGATGAGCGAAGAACAATTTGGCGGAGATGGTGCCGCAGCCGCAGC
TACAGCAGCAGTCGGTGGGTCTGCTGGAGAACAAGAAGGTGCAATGGTAGCC
GCAACACAAGGAGCGGCTGCAGCCGCAGGAAGTGGGGCAGGAACGGGCGGC
GGGACAGCCTCTGGCGGTACAGAGGGTGGATCTGCAGAAAGTGAAGGAGCC
AAAATAGATGCTTCTAAGAATGAAGAAGACGAGGGGCACAGCAATAGCTCA
CCTAGGCATAGCGAGGCCGCCACTGCGCAAAGAGAGGAGTGGAAGATGTTC
ATTGGTGGTCTCAGTTGGGATACCACCAAGAAGGACCTTAAAGATTATTTCA
GTAAGTTCGGCGAGGTGGTGGATTGTACCCTCAAACTGGACCCAATTACGGG
AAGAAGCCGGGGCTTCGGTTTCGTACTCTTCAAGGAGTCCGAAAGCGTCGAC
AAAGTGATGGACCAGAAGGAGCACAAGCTGAACGGTAAAGTCATCGACCCC
AAGCGAGCAAAGGCTATGAAGACTAAGGAACCTGTGAAGAAGATCTTCGTC
GGCGGGTTGAGCCCCGACACTCCCGAGGAAAAGATCCGAGAATATTTCGGCG
GGTTCGGCGAAGTTGAGTCAATTGAACTGCCTATGGATAATAAAACTAACAA
ACGCCGAGGCTTTTGTTTCATAACATTCAAAGAAGAGGAGCCTGTTAAGAAA
ATCATGGAGAAGAAGTATCATAACGTGGGGTTATCCAAGTGCGAGATTAAGG
TGGCAATGAGTAAAGAGCAGTACCAACAACAACAGCAATGGGGCAGCCGTG
GCGGTTTTGCCGGCAGGGCACGGGGTAGGGGCGGGGATCAACAAAGCGGAT
ATGGCAAAGTGTCTCGCCGGGGCGGCCACCAGAACAGTTATAAGCCTTATTG
AGGGCCC3′

*AUF1-LgBiT P40 cloning*. A synthetic human AUF1 P40 gene fragment was purchased from Twist Bioscience and inserted into *Plasmid C* using standard cloning techniques with KpnI and AsiSI restriction enzymes to produce a C-terminal LgBiT tag. Gene fragment contains a Kozak sequence at the N-terminus.

**Gene Fragment**:
5′GGTACCGCCACCATGAGCGAAGAACAATTTGGCGGAGATGGAGCAGCCGC
CGCCGCCACAGCAGCCGTGGGTGGGAGTGCCGGAGAACAAGAAGGTGCAAT
GGTCGCAGCCACCCAAGGCGCTGCAGCAGCCGCCGGTTCTGGTGCAGGCACA
GGCGGTGGGACTGCAAGCGGCGGAACTGAGGGTGGGTCCGCGGAAAGCGAA
GGTGCCAAAATCGATGCGAGCAAGAATGAAGAAGACGAGGGTCACTCCAAT
TCATCTCCCCGGCATAGCGAGGCCGCCACCGCCCAAAGAGAGGAGTGGAAG

ATGTTCATCGGCGGGCTGTCATGGGATACAACTAAGAAGGACCTCAAAGATT
ATTTCAGCAAGTTCGGGGAAGTAGTGGATTGTACGCTCAAACTGGACCCCAT
TACCGGACGCTCTAGAGGGTTCGGGTTCGTCCTCTTCAAGGAGTCCGAATCCG
TTGACAAAGTAATGGACCAGAAGGAGCACAAGCTCAACGGTAAAGTCATCG
ACCCAAAGCGCGCAAAGGCGATGAAGACCAAGGAACCTGTCAAGAAGATCT
TCGTGGGCGGGCTGAGTCCCGACACCCCCGAGGAAAAGATTAGAGAATATTT
CGGCGGATTCGGCGAAGTTGAGTCAATTGAACTGCCAATGGATAATAAAACA
AACAAACGCAGAGGATTTTGTTTCATCACATTCAAAGAGGAAGAGCCCGTAA
AGAAAATCATGGAGAAGAAGTATCATAACGTGGGATTGAGCAAGTGCGAGA
TTAAGGTGGCTATGAGTAAAGAGCAGTACCAACAGCAACAGCAATGGGGAT
CACGAGGCGGGTTCGCCGGTAGGGCGCGCGGCCGTGGCGGGGATCAACAAT
CCGGGTACGGTAAAGTCAGTCGGAGAGGCGGACACCAGAACTCTTATAAGCC
CTATGCGATCGC3′

*LgBiT-AUF1 P42 cloning*. A synthetic human AUF1 P42 gene fragment was purchased from Twist Bioscience and inserted into *Plasmid A* using standard cloning techniques with XhoI and ApaI restriction enzymes to produce an N-terminal LgBiT tag.

**Gene Fragment**:
5′CTCGAGATGAGCGAAGAACAATTTGGTGGGGATGGAGCCGCTGCTGCTGCC
ACTGCCGCCGTGGGCGGAAGTGCAGGAGAACAAGAAGGCGCAATGGTAGCC
GCAACACAAGGCGCGGCCGCCGCAGCTGGCTCAGGCGCTGGAACTGGAGGA
GGGACTGCATCCGGCGGGACAGAAGGCGGATCCGCGGAAAGCGAAGGTGCT
AAAATTGATGCTTCAAAGAATGAAGAAGACGAGGGTAAGATGTTCATTGGTG
GGTTGTCATGGGATACAACCAAGAAGGACCTCAAAGATTATTTCAGTAAGTT
CGGTGAGGTAGTTGATTGTACCCTTAAACTCGACCCAATTACGGGCCGGAGT
AGAGGCTTCGGGTTCGTTCTGTTCAAGGAGTCCGAATCTGTTGACAAAGTGAT
GGACCAGAAGGAGCACAAGCTGAACGGCAAAGTTATAGACCCGAAGCGGGC
TAAGGCAATGAAGACTAAGGAACCTGTGAAGAAGATATTCGTGGGCGGACTC
AGTCCGGACACCCCAGAGGAAAAGATTAGAGAATATTTCGGCGGATTCGGGG
AAGTCGAGAGCATCGAACTGCCTATGGATAATAAAACTAACAAACGCAGGG
GCTTTTGTTTCATCACATTCAAAGAAGAGGAGCCTGTCAAGAAAATCATGGA
GAAGAAGTATCATAACGTGGGGCTGAGCAAGTGCGAGATCAAGGTGGCAAT
GTCTAAAGAGCAGTACCAACAGCAACAGCAATGGGGTTCCCGGGGCGGCTTC
GCGGGTCGTGCCAGGGGCAGGGGTGGCGGACCGTCCCAGAATTGGAATCAA
GGCTACTCTAATTACTGGAACCAGGGTTACGGTAATTACGGCTACAATTCCCA
GGGATATGGCGGATACGGCGGCTACGATTATACGGGCTATAATAATTATTAC
GGCTACGGGGACTACAGTAATCAACAATCTGGCTACGGCAAAGTCAGCCGCA
GAGGTGGGCACCAGAACTCTTATAAGCCGTATTAGGGGCCC3′

*AUF1-LgBiT P42 cloning*. A synthetic human AUF1 P42 gene fragment was purchased from Twist Bioscience and inserted into *Plasmid C* using standard cloning techniques with KpnI and AsiSI restriction enzymes to produce a C-terminal LgBiT tag. Gene fragment contains a Kozak sequence at the N-terminus.

**Gene Fragment**:
5′GGTACCGCCACCATGAGCGAGGAACAATTCGGTGGAGATGGAGCAGCAGC
TGCTGCGACTGCAGCCGTTGGTGGTAGTGCCGGAGAACAAGAAGGCGCTATG
GTCGCTGCCACCCAAGGAGCTGCTGCAGCAGCCGGTTCTGGGGCAGGTACAG
GAGGAGGGACTGCATCAGGTGGGACAGAAGGCGGTTCAGCTGAAAGCGAAG
GCGCTAAAATCGATGCATCCAAGAATGAAGAAGACGAGGGTAAGATGTTCAT
TGGCGGGCTCTCTTGGGATACCACTAAGAAGGACCTCAAAGATTATTTCAGC
AAGTTCGGCGAGGTGGTGGATTGTACCCTCAAACTCGACCCGATTACCGGAA
GGTCCAGAGGGTTCGGATTCGTTCTCTTCAAGGAGAGCGAAAGCGTTGACAA
AGTGATGGACCAGAAGGAGCACAAGCTCAACGGTAAAGTAATCGACCCCAA
GCGGGCAAAGGCAATGAAGACCAAGGAACCTGTGAAGAAGATATTCGTGGG
CGGGCTGTCCCCGGACACCCCCGAGGAAAAGATTCGGGAATATTTCGGCGGC
TTCGGGGAAGTTGAGTCTATTGAACTGCCTATGGATAATAAAACAAACAAAC
GTCGCGGTTTCTGTTTCATAACATTCAAAGAGGAAGAGCCTGTTAAGAAGAT
TATGGAGAAGAAGTATCATAACGTGGGGCTGTCTAAGTGCGAGATCAAGGTG
GCTATGAGTAAAGAGCAGTATCAACAGCAACAGCAATGGGGTTCCCGCGGCG
GGTTCGCTGGCCGCGCACGCGGTCGAGGCGGCGGACCATCCCAGAATTGGAA
TCAAGGCTACTCCAATTACTGGAACCAGGGGTACGGTAATTACGGCTACAAT
TCACAGGGGTATGGCGGGTACGGCGGCTACGATTATACAGGGTATAATAATT
ATTACGGGTACGGGGACTACTCAAATCAACAAAGCGGATACGGCAAAGTGA
GCCGCAGAGGCGGGCACCAGAACTCTTATAAGCCTTATGCGATCGC3′

*LgBiT-AUF1 P45 cloning*. A synthetic human AUF1 P45 gene fragment was purchased from Twist Bioscience and inserted into *Plasmid A* using standard cloning techniques with XhoI and ApaI restriction enzymes to produce an N-terminal LgBiT tag.

**Gene Fragment**:
5′CTCGAGATGTCCGAGGAACAGTTCGGCGGGGACGGCGCAGCTGCAGCTGCC
ACAGCCGCCGTGGGGGGGTCCGCAGGAGAGCAGGAGGGGGCAATGGTTGCA
GCTACTCAGGGGGCCGCCGCTGCGGCTGGTTCAGGTGCCGGCACTGGTGGGG
GAACCGCCAGCGGAGGGACAGAGGGTGGGTCTGCAGAGAGCGAGGGGGCTA
AGATTGACGCTTCCAAAAATGAGGAAGATGAGGGGCATTCTAATTCATCCCC
ACGCCACTCCGAGGCCGCAACAGCACAGAGAGAGGAATGGAAGATGTTTAT
CGGAGGCCTCAGTTGGGATACAACGAAGAAGGATCTCAAGGACTACTTTTCA
AAGTTTGGCGAGGTAGTAGATTGTACACTGAAACTGGACCCAATCACAGGCC
GGAGTCGCGGCTTTGGATTTGTGCTGTTTAAGGAATCTGAGAGTGTGGATAA
AGTGATGGATCAGAAGGAGCATAAGCTTAATGGGAAAGTAATTGACCCCAA
GAGAGCAAAAGCTATGAAAACGAAAGAACCAGTCAAGAAAATCTTTGTCGG
GGGCCTGTCTCCTGACACTCCAGAAGAGAAGATCCGCGAATATTTTGGCGGA
TTCGGAGAAGTTGAAAGCATCGAGCTCCCAATGGACAACAAAACCAACAAG
CGCCGTGGCTTCTGCTTTATCACATTTAAGGAAGAGGAGCCGGTGAAGAAGA
TTATGGAGAAGAAATACCACAACGTGGGGTTGTCCAAGTGCGAGATAAAGGT
TGCTATGAGTAAGGAGCAGTACCAACAACAGCAACAGTGGGGATCAAGAGG
GGGGTTCGCCGGGCGTGCCCGGGGCCGAGGGGGAGGCCCATCCCAAAATTG
GAACCAGGGATACTCCAACTATTGGAACCAGGGTTATGGGAATTACGGGTAT

AATTCCCAGGGATATGGAGGCTATGGCGGCTATGATTATACGGGTTACAACA
ACTACTACGGCTATGGGGACTATTCCAATCAGCAGAGTGGCTATGGAAAGGT
CAGCCGCAGAGGGGGCCATCAAAACTCCTACAAGCCTTATTAAGGGCCC3′

*AUF1-LgBiT P45 cloning*. A synthetic human AUF1 P45 gene fragment was purchased from Twist Bioscience and inserted into *Plasmid C* using standard cloning techniques with KpnI and AsiSI restriction enzymes to produce a C-terminal LgBiT tag. Gene fragment contains a Kozak sequence at the N-terminus.

**Gene Fragment**:
5′GGTACCGCCACCATGTCCGAGGAACAGTTCGGCGGGGACGGCGCAGCTGC
AGCTGCCACAGCCGCCGTGGGGGGGTCCGCAGGAGAGCAGGAGGGGGCAAT
GGTTGCAGCTACTCAGGGGGCCGCCGCTGCGGCTGGTTCAGGTGCCGGCACT
GGTGGGGGAACCGCCAGCGGAGGGACAGAGGGTGGGTCTGCAGAGAGCGAG
GGGGCTAAGATTGACGCTTCCAAAAATGAGGAAGATGAGGGGCATTCTAATT
CATCCCCACGCCACTCCGAGGCCGCAACAGCACAGAGAGAGGAATGGAAGA
TGTTTATCGGAGGCCTCAGTTGGGATACAACGAAGAAGGATCTCAAGGACTA
CTTTTCAAAGTTTGGCGAGGTAGTAGATTGTACACTGAAACTGGACCCAATC
ACAGGCCGGAGTCGCGGCTTTGGATTTGTGCTGTTTAAGGAATCTGAGAGTG
TGGATAAAGTGATGGATCAGAAGGAGCATAAGCTTAATGGGAAGTAATTG
ACCCCAAGAGAGCAAAAGCTATGAAAACGAAAGAACCAGTCAAGAAAATCT
TTGTCGGGGGCCTGTCTCCTGACACTCCAGAAGAGAAGATCCGCGAATATTTT
GGCGGATTCGGAGAAGTTGAAAGCATCGAGCTCCCAATGGACAACAAAACC
AACAAGCGCCGTGGCTTCTGCTTTATCACATTTAAGGAAGAGGAGCCGGTGA
AGAAGATTATGGAGAAGAAATACCACAACGTGGGGTTGTCCAAGTGCGAGA
TAAAGGTTGCTATGAGTAAGGAGCAGTACCAACAACAGCAACAGTGGGGAT
CAAGAGGGGGGTTCGCCGGGCGTGCCCGGGGCCGAGGGGGAGGCCCATCCC
AAAATTGGAACCAGGGATACTCCAACTATTGGAACCAGGGTTATGGGAATTA
CGGGTATAATTCCCAGGGATATGGAGGCTATGGCGGCTATGATTATACGGGT
TACAACAACTACTACGGCTATGGGGACTATTCCAATCAGCAGAGTGGCTATG
GAAAGGTCAGCCGCAGAGGGGGCCATCAAAACTCCTACAAGCCTTATGCGAT
CGC3′

### B. RNA Probes

**Table S1.** Sequences of RNA probes used in this study.

| RNA | Sequence |
|---|---|
| **(CUG)$_{19}$ U16** | CGCUGCUGCUGCUGC(**5-LC-N-U**)GCUGCUGCUGCUGCUGCUGCUGCUGC UGCUGCUGCUGCUGCG |
| **(CUG)$_{19}$ U31** | CGCUGCUGCUGCUGCUGCUGCUGCUGC(**5-LC-N-U**)GCUGCUGCUGCUGC UGCUGCUGCUGCUGCG |
| **(GUC)$_{19}$ U16** | GCGUCGUCGUCGUCG(**5-LC-N-U**)CGUCGUCGUCGUCGUCGUCGUCGUCG UCGUCGUCGUCGUCGC |
| **(GUC)$_{19}$ U31** | GCGUCGUCGUCGUCGUCGUCGUCGUCG(**5-LC-N-U**)CGUCGUCGUCGUCG UCGUCGUCGUCGUCGC |
| **Numb1 U33** | GCACACUGCGAAAUGGUCAUACCGAUGACCCG(**5-LC-N-U**)GCUCACUAGGUA GUAGUUUUAAUAAGUGAGCAAGCAGUGUUGUCCU |
| **Numb1 U52** | GCACACUGCGAAAUGGUCAUACCGAUGACCCGUGCUCACUAGGUAGUAGUU (**5-LC-N-U**)UAAUAAGUGAGCAAGCAGUGUUGUCCU |
| **Numb1 U56** | GCACACUGCGAAAUGGUCAUACCGAUGACCCGUGCUCACUAGGUAGUAGUU UUAA(**5-LC-N-U**)AAGUGAGCAAGCAGUGUUGUCCU |
| **Numb1 Mut. U33** | GCACACUGCGAAAUGGUCAUACCGAUGACCCG(**5-LC-N-U**)GCUCACUAGGAA GAAGUUUUAAUAAGUGAGCAAGCAGUGUUGUCCU |
| **Numb1 Mut. U52** | GCACACUGCGAAAUGGUCAUACCGAUGACCCGUGCUCACUAGGAAGAAGUU (**5-LC-N-U**)UAAUAAGUGAGCAAGCAGUGUUGUCCU |
| **Numb1 Mut. U56** | GCACACUGCGAAAUGGUCAUACCGAUGACCCGUGCUCACUAGGAAGAAGUU UUAA(**5-LC-N-U**)AAGUGAGCAAGCAGUGUUGUCCU |
| **cfos ARE U2** | U(**5-LC-N-U**)UUAUUGUUUUAAUUUAUUUAUUAAGAUGGAUUCUCAGAUAUU UA |
| **cfos ARE U23** | UUUUAUUGUUUUAAUUUAUUUA(**5-LC-N-U**)UAAGAUGGAUUCUCAGAUAUU UA |
| **cfos ARE U36** | UUUUAUUGUUUUAAUUUAUUUAUUAAGAUGGAUUC(**5-LC-N-U**)CAGAUAUU UA |
| **cfos ARE 1-26 U2** | U(**5-LC-N-U**)UUAUUGUUUUAAUUUAUUUAUUAA |
| **cfos ARE 1-26 U23** | UUUUAUUGUUUUAAUUUAUUUA(**5-LC-N-U**)UAA |
| **cfos ARE 13-37 U23** | AAUUUAUUUA(**5-LC-N-U**)UAAGAUGGAUUCUC |
| **cfos ARE 13-37 U36** | AAUUUAUUUAUUAAGAUGGAUUC(**5-LC-N-U**)C |
| **Rβ31 U8** | UGGCCAA(**5-LC-N-U**)GCCCUGGCUCACAAAUACCACUG |
| **Rβ31 U17** | UGGCCAAUGCCCUGGC(**5-LC-N-U**)CACAAAUACCACUG |

All RNA sequences contain a 5′ biotin modification with an 18-atom spacer. The HaloTag® ligand was conjugated to the amine of the 5-aminohexylacrylamino-uridine (5-LC-N-U) modification as described in the Materials and Methods.

**Table S2.** Chemical structures of the modified uridine base used in RNA substrates for RiPCA (5-LC-N-U), the HaloTag® ligand used to modify 5-LC-N-U-containing RNAs, and the product of the labeling reaction.

| Name | Structure |
| --- | --- |
| 5-aminohexylacrylamino-uridine (5-LC-N-U) | 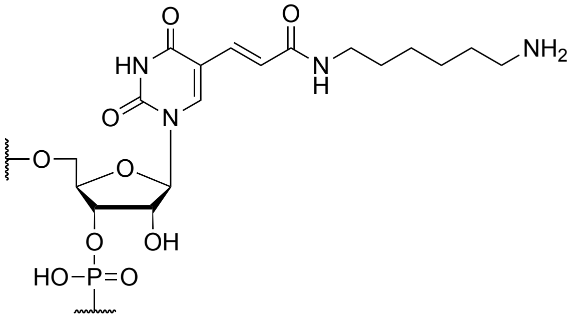 |
| HaloTag Succinimidyl Ester (O2) Ligand | 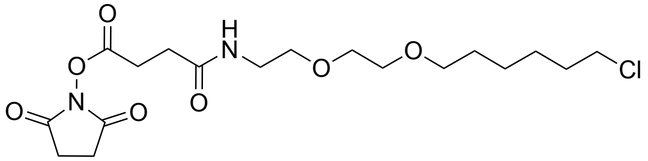 |
| 5-LC-N-U Base conjugated to HaloTag Ligand | 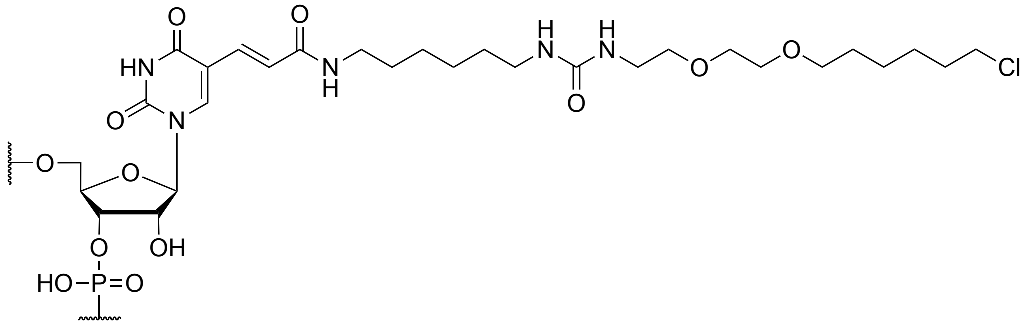 |

**Table S3.** Affinities for RNA-protein interactions^ investigated in this study.

| RBP | RNA Substrate | Affinity | Method |
| --- | --- | --- | --- |
| ***MBNL1*** | $(CUG)_4$ | 3.1 ± 0.1 nM[2] | *TIRFM binding assay |
| | $(CUG)_{54}$ | 5.3 ± 0.6 nM[3] | Filter binding assay |
| ***Stau1*** | GC-rich RNA with 2° structure | ~17-30 nM[4] | Fluorescence anisotropy (FA) |
| ***CELF1*** | $(CUG)_{10}$ | 27 ± 12 μM[5] | **SPR |
| | $(CUG)_{140}$ | 6 ± 2 μM[5] | **SPR |
| ***Msi*** | 5' UAGUAG 3' sequence | ~4 nM[6] | Gel retardation assay |
| ***AUF1 P37*** | *c-fos* RNA substrate | 7.8 nM[7] | ***Quantitative EMSA |

^These RNA-RBP interactions represent the closest comparison to the ones used in this work which have reported binding affinities in the literature. *TIRFM = Total Internal Reflection Fluorescence Microscope. **SPR = Surface Plasmon Resonance. ***EMSA = Electrophoretic Mobility Shift Assay.

### C. Supplemental Figures

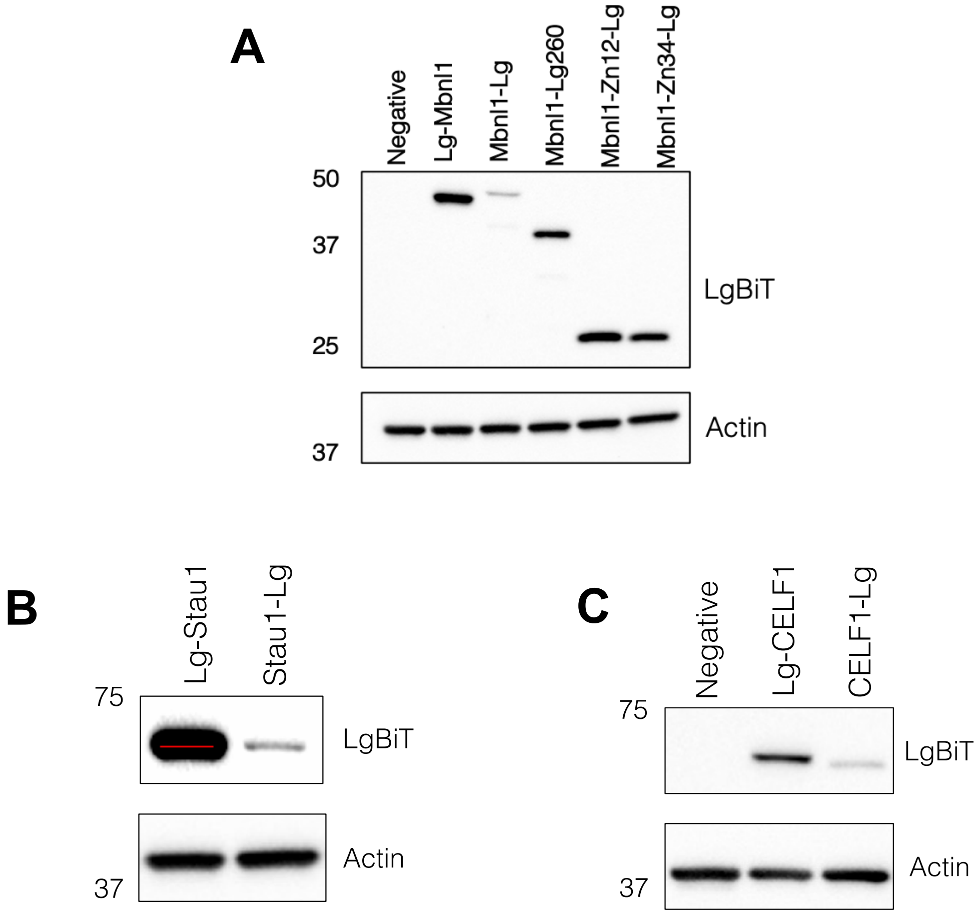


**Figure S1**. Expression of LgBiT-tagged RBPs used in RiPCA in Flp-In 293 cells stably expressing SmBiT-HaloTag. (A) MBNL1 constructs. (B) Stau1. (C) CELF1.

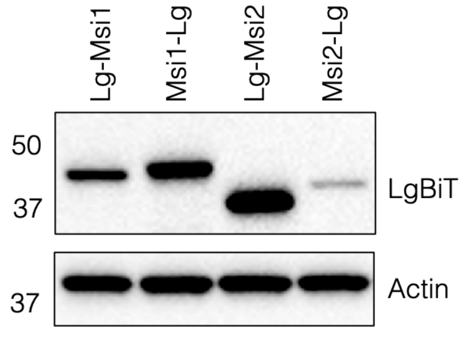


**Figure S2**. Expression of LgBiT-tagged Msi1 and Msi2 in Flp-In 293 cells stably expressing SmBiT-HaloTag.

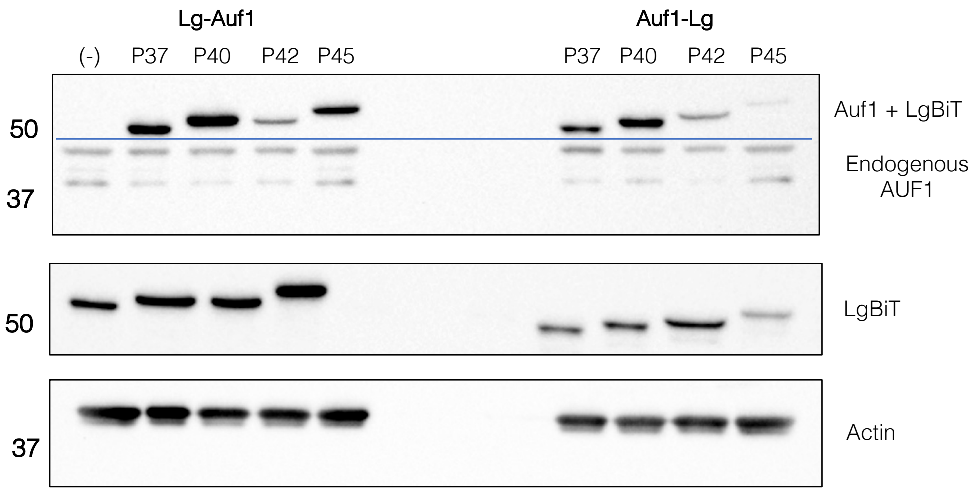


**Figure S3**. Expression of LgBiT-tagged Auf1 isoforms in Flp-In 293 cells stably expressing SmBiT-HaloTag and endogenous Auf1 levels.

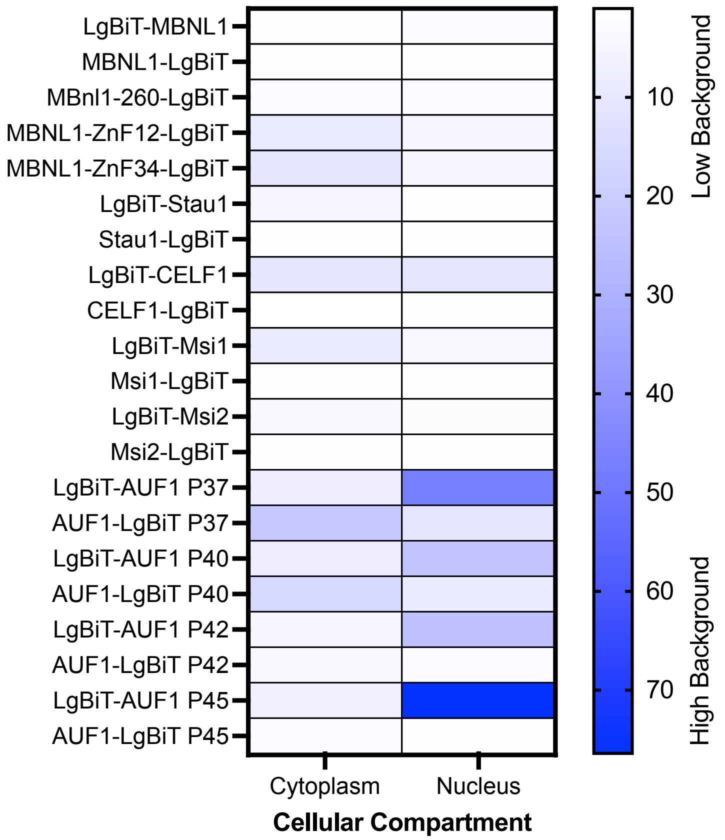


**Figure S4**. Heat map depicting the background values obtained with each of the protein constructs in the nuclear and cytoplasmic compartments using 1.0 ng/well of plasmid. Background values were obtained by dividing the measured chemiluminescence values from assays transfected with each of the plasmids but no RNA by the measured chemiluminescence values from assays not transfected with an RBP-LgBiT plasmid or RNA (cells only). Cytoplasmic values were obtained using the SmBiT-HaloTag cell line and nuclear values were obtained using the SmBiT-HaloTag-NLS cell line.

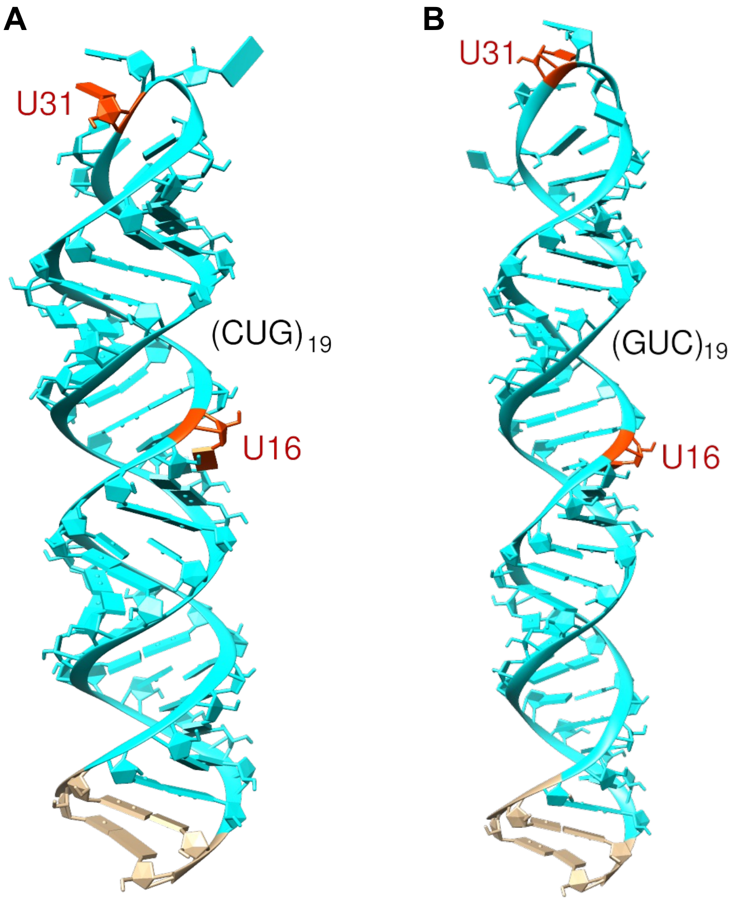


**Figure S5.** 3D Structure Prediction of RNA substrates for repeat RNA systems. Cyan: RBP binding site, Red: U modification site. (A) $(CUG)_{19}$ probe. (B) $(GUC)_{19}$ probe. [8, 9]

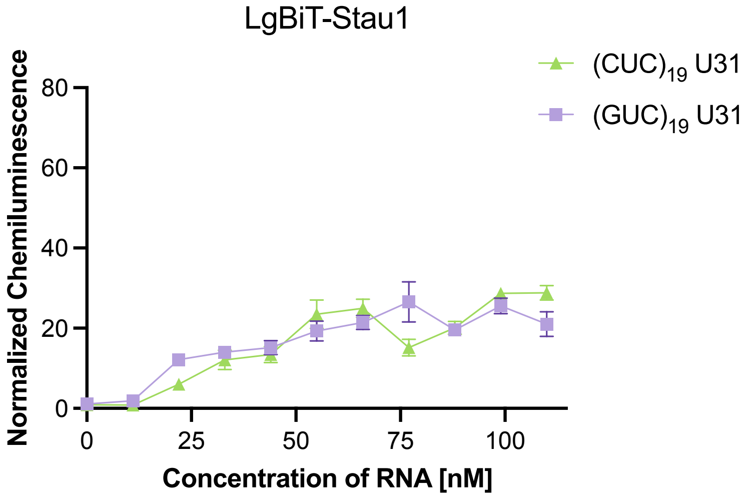


**Figure S6.** Dose-dependence of $(CUG)_n$ expanded repeat RiPCA with Stau1 as compared to $(GUC)_n$ sequence. Normalized chemiluminescence values obtained using (1.0ng/well) LgBiT-Stau1.

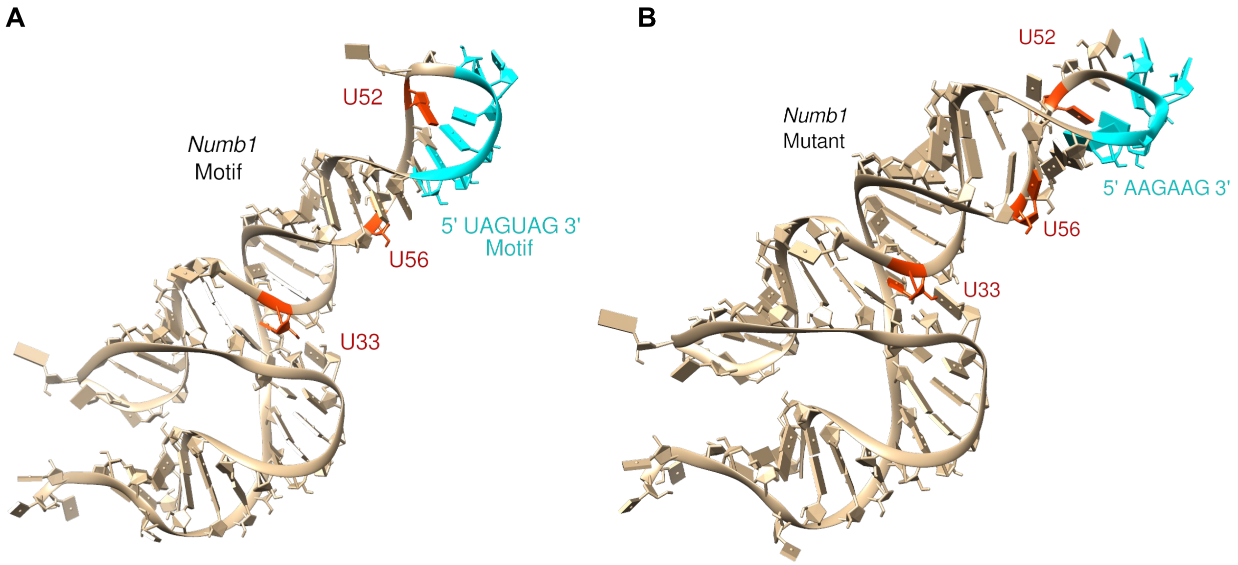


**Figure S7.** 3D Structure Prediction of RNA substrates tested with Msi1/2. Cyan: RBP binding site (or mutated RBP binding site), Red: U modification site. (A) *Numb1* RNA. (B) Mutant *Numb1* RNA.[8, 9]

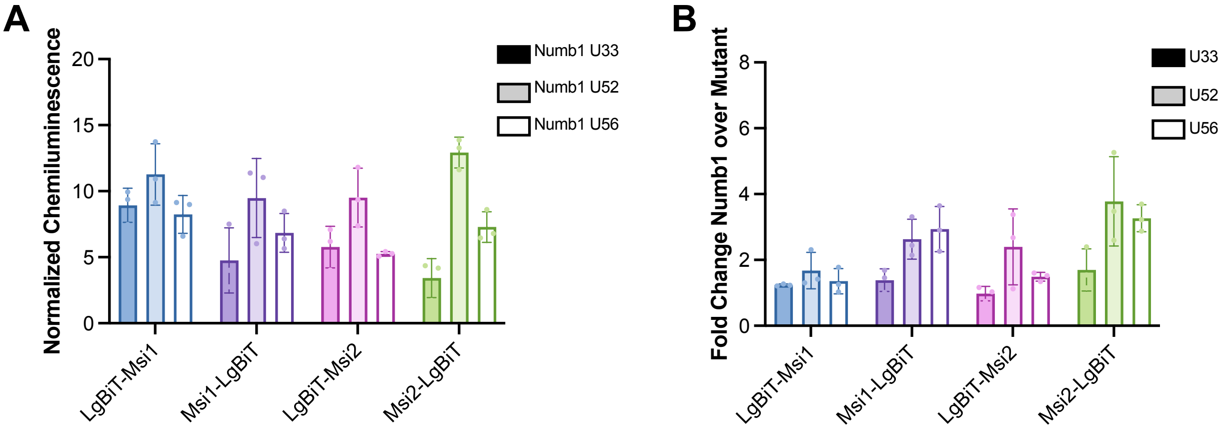


**Figure S8.** *Numb1* RNA motif with Msi1 and Msi2. (A) Normalized chemiluminescence values calculated from nuclear assays. (B) Binding selectivity for *numb1* RNAs over non-binding mutant *numb1* RNAs at each of the modification sites detected in the nucleus.

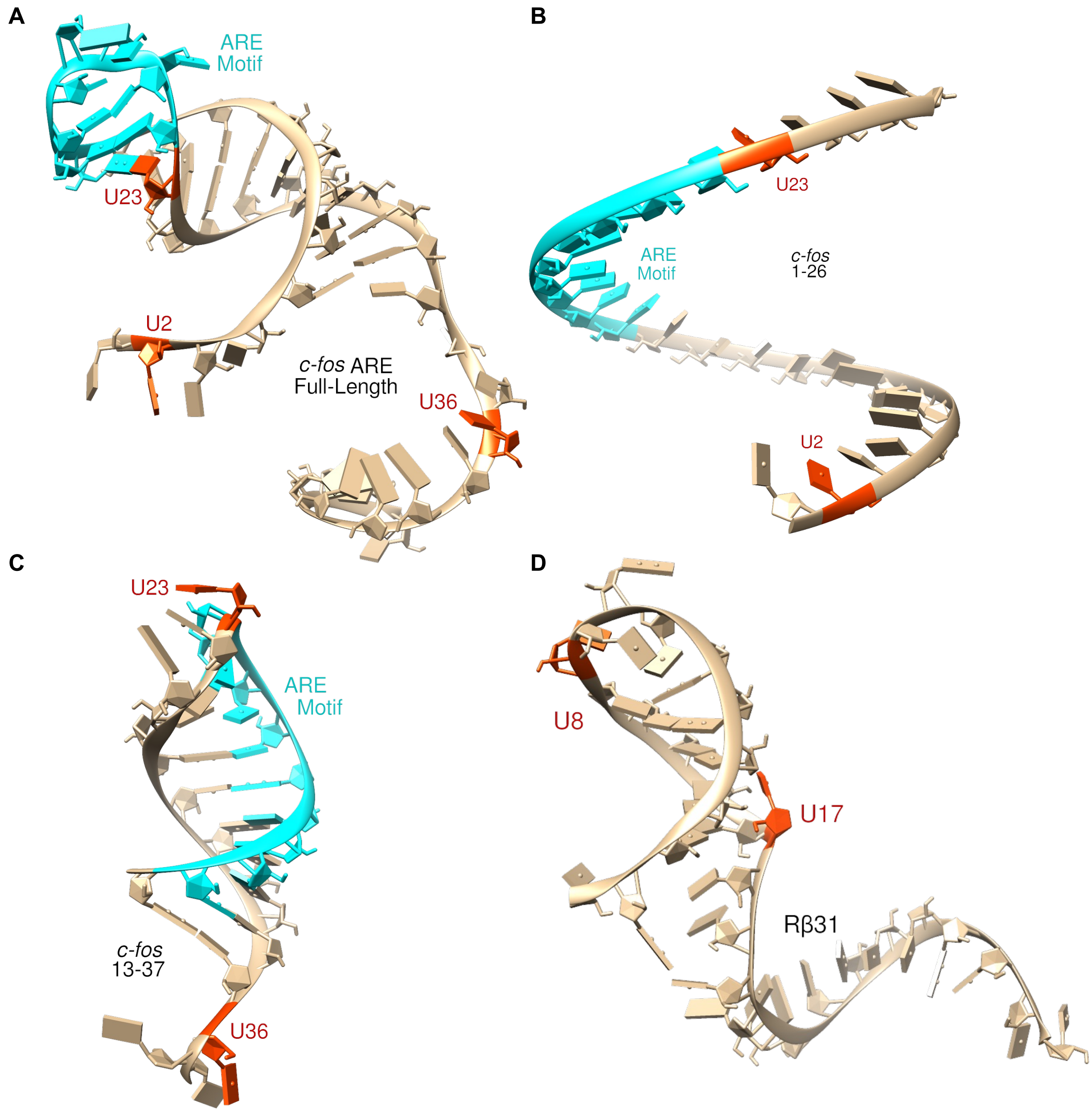


**Figure S9.** 3D Structure Prediction of RNA substrates tested with AUF1 proteins. Cyan: RBP binding site, Red: U modification site. (A) Full-length *c-fos* ARE. (B) *c-fos* 1-26. (C) *c-fos* 13-37. (D) Rβ31.[8, 9]

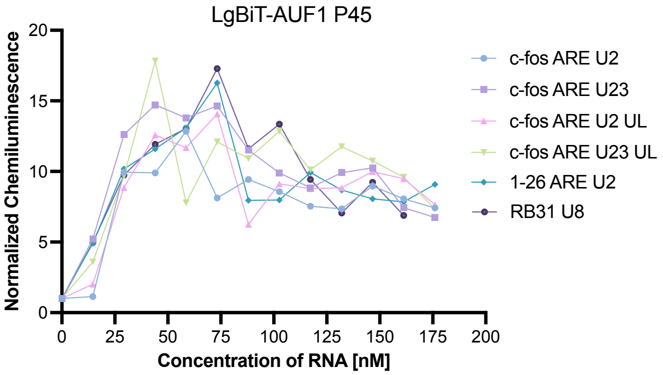


**Figure S10.** Dose-dependence of various substrates in RiPCA with LgBiT-AUF1 P45 (1.2 ng/well) including positive binders (c-fos ARE U2, c-fos ARE U23, and 1-26 ARE U2) and negative binders (c-fos ARE U2 UL, c-fos ARE U23 UL, Rβ31 U8). UL = unlabeled RNA and indicates that the RNA is not conjugated to the HaloTag® ligand and cannot be labeled by SmBiT-HaloTag in cells.

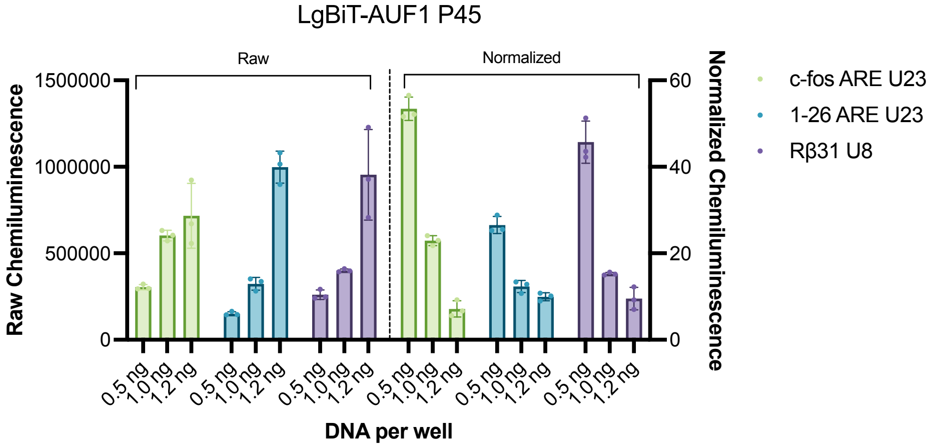


**Figure S11.** Raw and normalized chemiluminescence values obtained with LgBiT-AUF1 P45 and select RNA substrates from Figure 3F with increasing amounts of LgBiT-AUF1 P45. LgBiT-RBP = N-terminally tagged RBP. RBP-LgBiT = C-terminally tagged RBP.

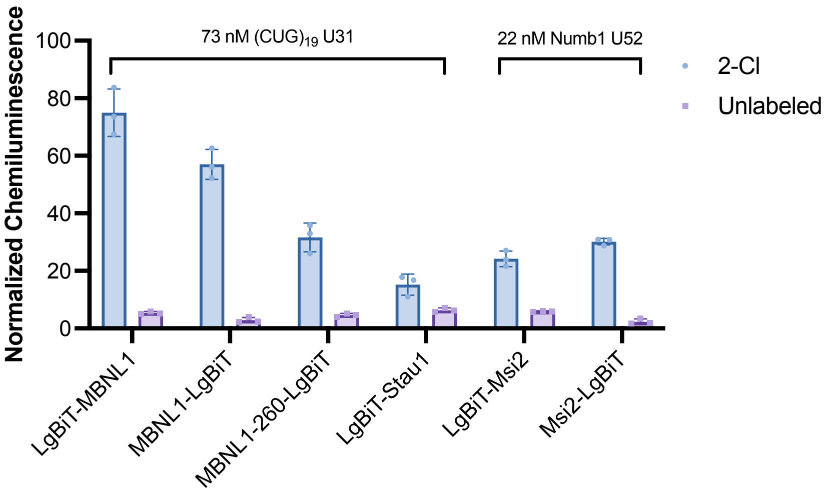


**Figure S12**. Normalized chemiluminescence values obtained with RBPs and their HaloTag® ligand-labeled and unlabeled RNAs.

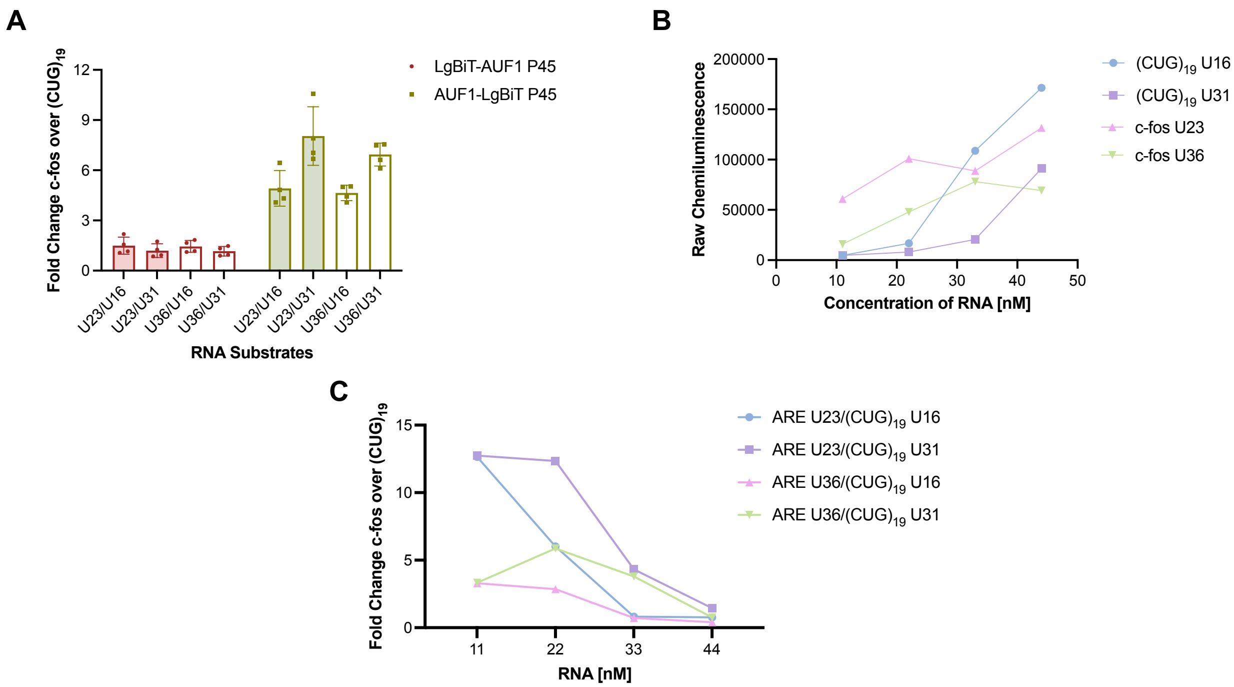


**Figure S13**. Specificity of LgBiT-AUF1 P45 and AUF1-LgBiT P45 in comparison to $(CUG)_n$ RNAs. (A) Binding selectivity for LgBiT-AUF1 P45 and AUF1-LgBiT P45 for *c-fos* ARE U23 and U36 binding over $(CUG)_{19}$ U16 and U31. Values calculated using 1.0 ng/well of each plasmid and 33 nM of each RNA in the cytoplasmic assay. (B) Raw chemiluminescence values of LgBiT-Auf1 P45 (0.5 ng/well) tested with *c-fos* ARE (positive substrate) and $(CUG)_{19}$ RNA as a non-binding control. (C) Data from S12B plotted as the signal generated with LgBiT-AUF1 P45 and *c-fos* ARE divided by the signal generated by LgBiT-AUF1 P45 with $(CUG)_{19}$ RNAs. Binding specificity of LgBiT-AUF1 P45 for *c-fos* ARE RNA over $(CUG)_{19}$ RNA demonstrated only at low concentrations of the RNA.

### D. Supplemental Tables

**Table S4.** RiPCA conditions.

| Component | Amount Added (triplicate conditions) | Final Amounts/Concentrations per well |
|---|---|---|
| OptiMEM | 37.5 µL | N/A |
| 3.9 ng/µL DNA | 0.866 µL | ~1 ng |
| 25 µM RNA | 0.45 µL | ~33 nM |
| TransIT-X2 | 1.126 µL | N/A |
| 200,000 cells/mL | 300 µL | 20,000 cells/well |
| **Total Final Volume of Mixture** | **~340 µL** | **100 µL per well** |
